# PTK7 is a positive allosteric modulator of GPR133 (ADGRD1) signaling in GBM

**DOI:** 10.1101/2022.06.15.496232

**Authors:** Joshua D. Frenster, Hediye Erdjument-Bromage, Wenke Liu, Gabriele Stephan, Niklas Ravn-Boess, Devin Bready, Jordan Wilcox, Björn Kieslich, Manuel Jankovic, Caroline Wilde, Susanne Horn, Norbert Sträter, Ines Liebscher, Torsten Schöneberg, David Fenyo, Thomas A. Neubert, Dimitris G. Placantonakis

## Abstract

GPR133 (ADGRD1), an adhesion G protein-coupled receptor, supports growth of glioblastoma, a brain malignancy. We demonstrated that GPR133 is intramolecularly cleaved, and that dissociation of its N-terminal and C-terminal fragments (NTF and CTF) at the plasma membrane correlates with increased receptor signaling. However, how the extracellular interactome of GPR133 in glioblastoma modulates signaling remains unknown. Here, we use affinity purification and mass spectrometry to identify extracellular binding partners of GPR133 in patient-derived glioblastoma cells. We show that the transmembrane protein PTK7 binds the GPR133 NTF and its expression *in trans* increases GPR133 signaling. This effect requires the intramolecular cleavage of GPR133 and PTK7’s anchoring in the plasma membrane. The GPR133-PTK7 interaction facilitates orthosteric activation of GPR133 by soluble peptide mimicking the endogenous tethered *Stachel* agonist, suggesting PTK7 binding allosterically enhances accessibility of GPR133’s orthosteric *Stachel* binding pocket. GPR133 and PTK7 are expressed in adjacent cells in glioblastoma, where their knockdown phenocopies each other. We propose that this novel ligand-receptor interaction is relevant to the pathogenesis of glioblastoma, as well as physiological processes in several tissues.

## INTRODUCTION

Adhesion G protein-coupled receptors (aGPCRs) are thought to serve dual functions as both transmembrane adhesion proteins and transducers of extracellular signals. This protein class is characterized by their long extracellular N-termini, which contain receptor-specific adhesive domains that help define the extracellular interactome (Langenhan *et al*, 2013; Vizurraga *et al*, 2020), as well as the phylogenetically conserved GPCR autoproteolysis-inducing (GAIN) domain that catalyzes intramolecular cleavage of the N-terminus at the GPCR proteolysis site (GPS) (Arac *et al*, 2012). In C-terminal proximity to the GPS lies an endogenous tethered agonist, the *Stachel* sequence, that is essential to receptor activation (Barros-Alvarez *et al*, 2022; Liebscher *et al*, 2014; Ping *et al*, 2022; Qu *et al*, 2022; Xiao *et al*, 2022). A popular model for aGPCR activation invokes ligand binding as a means to convey mechanical forces driving the dissociation of the cleaved extracellular N-terminal fragment (NTF) from the remaining transmembrane-spanning C-terminal fragment (CTF), which contains the *Stachel* sequence (Lin *et al*, 2022; Wilde *et al*, 2022). This dissociation, in turn, is thought to allow the *Stachel* sequence to insert itself into the orthosteric binding pocket formed by the transmembrane portion of the receptor to exert agonistic effects (Barros-Alvarez *et al*., 2022; Ping *et al*., 2022; Qu *et al*., 2022; Xiao *et al*., 2022). In an alternative model inspired by the fact that cleavage-deficient mutant aGPCRs maintain signaling competency, ligand binding and mechanical forces allosterically induce conformational changes leading to receptor activation and signaling dependent on orthosteric *Stachel* agonism but without the requirement for NTF-CTF dissociation. Both these models gained support through the recent publication of cryo-EM structures of several aGPCRs in their activated state identifying critical residues mediating the interaction of the *Stachel* sequence with its orthosteric binding pocket within the transmembrane portion of the CTF (Barros-Alvarez *et al*., 2022; Ping *et al*., 2022; Qu *et al*., 2022; Xiao *et al*., 2022).

GPR133 (ADGRD1) is a member of the aGPCR class, whose signaling requires the intramolecular agonistic *Stachel* sequence and leads to Gs-mediated increases in cytosolic Camp (Barros-Alvarez *et al*., 2022; Bohnekamp & Schoneberg, 2011; Liebscher *et al*., 2014; Ping *et al*., 2022; Qu *et al*., 2022; Xiao *et al*., 2022). Elucidating GPR133’s mechanisms of action and activation is particularly relevant in the context of glioblastoma (GBM), a primary brain malignancy, in which we found GPR133 to be expressed *de novo* relative to healthy brain and to be required for tumor growth (Bayin *et al*, 2016; Frenster *et al*, 2020). We recently demonstrated that GPR133 undergoes autoproteolytic cleavage at the GPS almost immediately after protein synthesis and that the NTF and CTF remain non-covalently bound in the secretory pathway, until they reach the plasma membrane, where dissociation occurs (Frenster *et al*, 2021). The cleavage and NTF-CTF of GPR133 correlate with increased receptor signaling, and cleavage is required for positive allosteric modulation of signaling induced by treatment with antibodies targeting the N-terminus (Frenster *et al*., 2021; Stephan *et al*, 2022). These findings invite a model in which the extracellular interactome of GPR133 and its effects on NTF-CTF dissociation and receptor conformation are predicted to allosterically modulate receptor signaling. However, the identity of extracellular interactors of GPR133 in GBM, as well as whether mechanical forces generated by their binding are involved in receptor signaling, remain unknown. The only previously known GPR133 ligand, the transmembrane cell surface protein PLXDC2 (Plexin Domain Containing 2), was recently shown to mediate an interaction with GPR133 critical to the female reproductive system in mice (Bianchi *et al*, 2021). However, the underlying mechanism of this interaction, including effects on NTF-CTF dissociation and mechano-activation requirements, were not assessed.

In this study, we used affinity co-purification and mass spectrometry to discover a number of candidate extracellular binding partners of GPR133 in patient-derived GBM cells. Among them, the single transmembrane-span protein tyrosine kinase 7 (PTK7) binds and activates GPR133 signaling robustly, when the receptor and the ligand are expressed in adjacent cells *in trans*. The allosteric activating effect of PTK7 on receptor signaling requires GPR133 cleavage and PTK7 anchoring to cell membranes or rigid extracellular substrates. Furthermore, the PTK7-GPR133 interaction synergizes with soluble *Stachel* peptide treatment to produce supralinear receptor activation. We show that PTK7 and GPR133 are both expressed in GBM specimens, often in complementary patterns, and that PTK7 knockdown phenocopies GPR133 knockdown in patient-derived GBM cells. These findings support a model in which PTK7 binding *in trans* allosterically increases GPR133 activation and signaling.

## RESULTS

### The extracellular interactome of GPR133

To identify the extracellular and plasma membrane-bound interactome of GPR133 in GBM, we utilized an affinity co-purification/mass spectrometry approach in patient-derived GBM cell cultures (Fig. 1A). Firstly, we constructed a GPR133 bait consisting of the full-length receptor with an intracellular (C-terminal) TwinStrep affinity tag and the H543R mutation which prevents autoproteolytic cleavage (Arac *et al*., 2012; Frenster *et al*., 2021; Krasnoperov *et al*, 2002; Lin *et al*, 2010; Moriguchi *et al*, 2004; Okajima *et al*, 2010; Volynski *et al*, 2004). This mutation was added to prevent the dissociation of the NTF, and thus allowed for the co-purification of extracellular interactors via the C-terminal affinity tag. We transduced three separate patient-derived GBM cultures with the lentiviral expression construct for this bait or a control empty vector, let them grow to confluence to increase cell-cell contacts, crosslinked the extracellular interactome with the linker DTSSP (3,3′-dithiobis(sulfosuccinimidyl propionate)) and purified GPR133 and its interacting proteins using StrepTactin in three biological repeats for each condition (Fig. 1B; Fig. EV1A). Mass spectrometric analysis of the eluates detected 2,138 proteins, of which 569 were either extracellular or membrane-bound (Fig. 1C). Of these, 87 proteins with biologically relevant subcellular compartment annotation showed enrichment within the GPR133 bait condition in all three patient-derived cell cultures, and 23 of them reached statistical significance (p<0.05) (Fig. 1D, E; Figure EV1B). We found that, among the significantly enriched (p<0.05) protein pool prior to filtering for appropriate subcellular localization (n=55), membrane proteins were overrepresented when compared to the total protein pool, which instilled confidence in our approach (Fig. EV1C).

**Figure 1.**
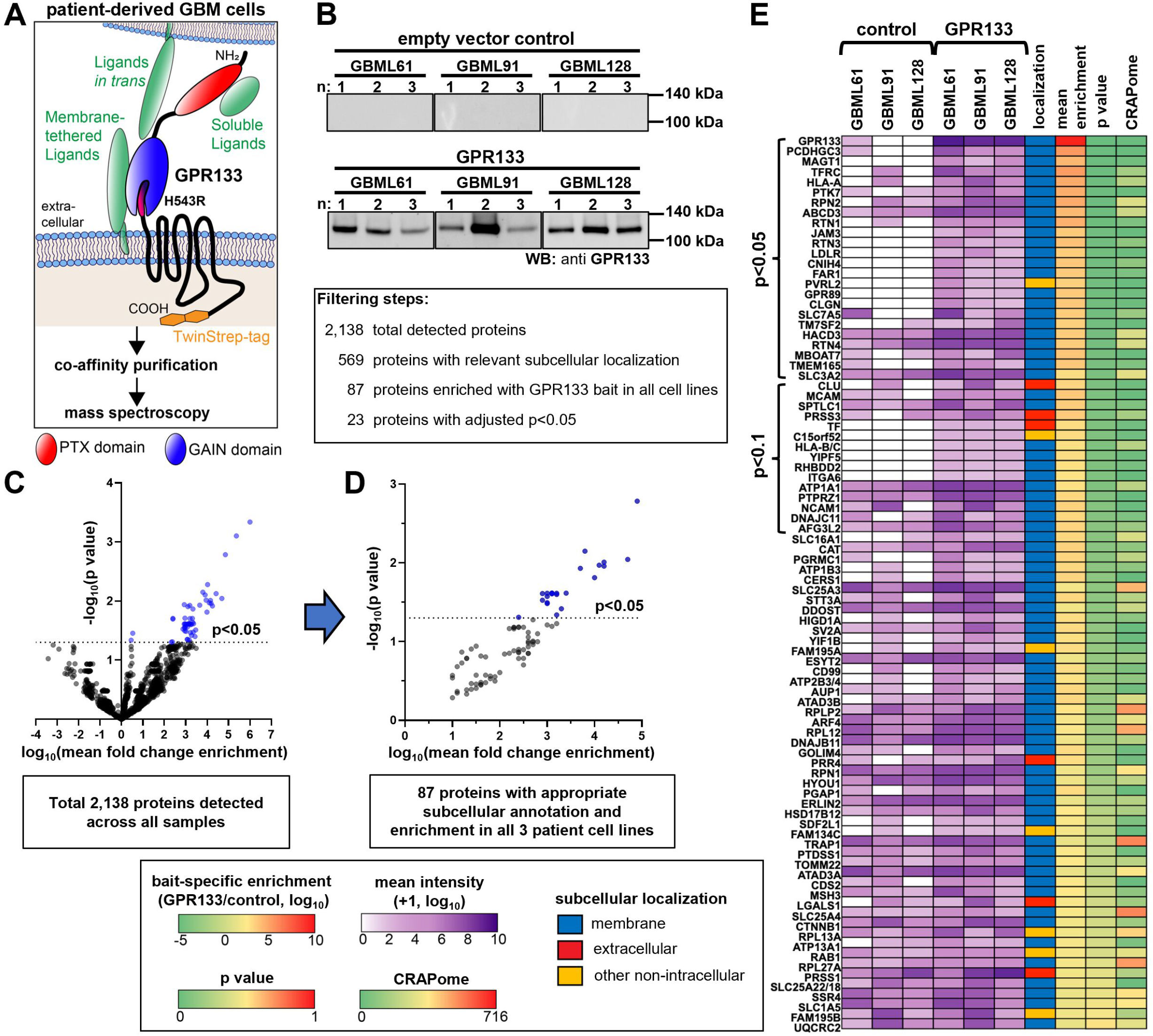
Co-purification/mass spectrometry-based ligand discovery in patient-derived GBM cells detects GPR133’s extracellular and membrane-bound interactome. A) Schematic overview of the affinity co-purification approach. Full-length cleavage-deficient GPR133-H543R with a C-terminal intracellular TwinStrep tag was overexpressed in patient-derived GBM cultures. GPR133 and binding partners were crosslinked with DTSSP, followed by affinity enrichment and analysis by mass spectrometry. B) Eluates after StrepTactin purification from 3 different GBM cultures (with 3 independent biological repeats each) transduced with either TwinStrep-tagged uncleavable GPR133-H543R or an empty vector were analyzed with SDS-PAGE and Western blot for GPR133 levels using an anti-GPR133 C-terminus antibody. C) Volcano plot of fold enrichment with the GPR133 bait and statistical significance of 2,138 total detected proteins. The 569 candidate proteins with subcellular annotation at the plasma membrane or in the extracellular space were filtered for enrichment with the GPR133 bait versus control vector samples across all cell lines, resulting in 87 consistently enriched proteins. D) Volcano plot depicting these 87 proteins, 23 of which were significantly enriched (p<0.05; blue dots). E) Overview heatmap of the 87 consistently enriched GPR133 binding protein candidates with subcellular localizations at the plasma membrane or extracellular space. The purple heatmap depicts detected protein intensities as a log_10_ mean of three biological replicates per culture per condition. The subcellular localization column was derived using Ensembl database annotations and is detailed in the Methods section. The mean enrichment column depicts the ratio of log_10_ mean intensities across all cell cultures and replicates from the GPR133 bait samples over the empty vector control samples. The p value column depicts the p value calculated from a mixed effects model of enrichment with the GPR133 bait across all samples. The CRAPome column depicts the frequency of detection of proteins by mass spectrometry across 716 publicly available affinity purification datasets, and thereby identifies possibly non-specific interactors.

### PTK7 binds GPR133

To validate binding between GPR133 and some of the top interactors identified in our screen, we generated HEK293T cell lines stably overexpressing TwinStrep-tagged uncleavable GPR133-H543R or an empty vector, and transfected them with Myc-tagged candidate proteins (PCDHGC3, PTK7, TFRC, NECTIN2, CADM4, MCAM) (Fig. 2Ai). Among these candidates PTK7, and to a lesser extent TFRC (transferrin receptor), demonstrated the most prominent binding to GPR133 in StrepTactin-mediated affinity co-purification assays (Fig. 2Aii, n=3 independent experiments). PTK7 is a type I single-pass transmembrane protein consisting of seven extracellular Ig-like C2-type repeats and an intracellular inactive tyrosine kinase domain (Fig. 2B), and has been previously implicated in the pathogenesis of GBM (Liu *et al*, 2015). PTK7 not only ranked among the top GPR133 bait-enriched proteins in our screen (Fig. 2C), but was also highlighted by the reproducibility of its co-purified protein intensity in 8 out of 9 biological replicates when normalized to the GPR133 bait (Fig. 2D; Fig. EV2).

**Figure 2.**
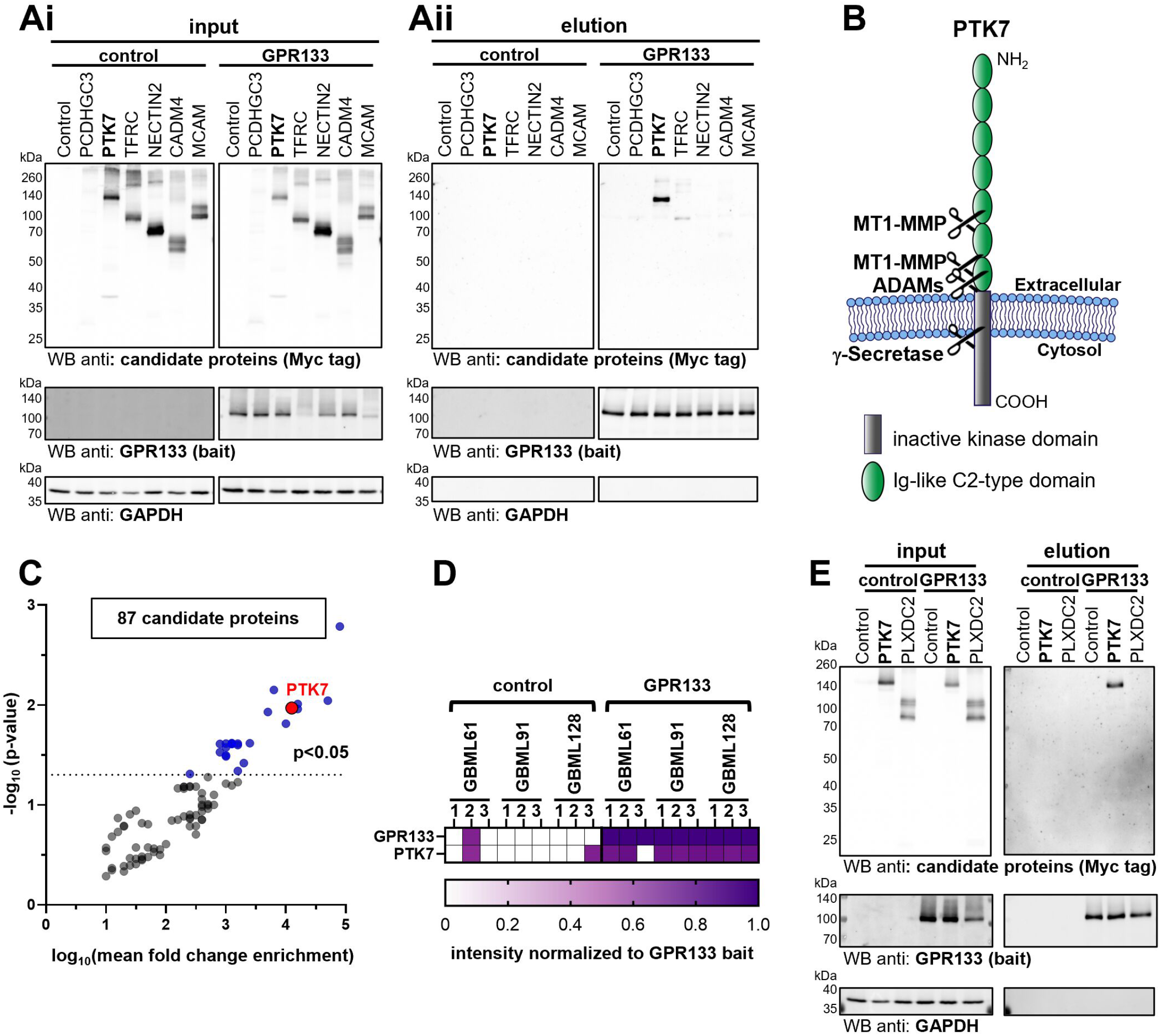
Biochemical validation of candidate binding proteins detects PTK7 as the most robust GPR133 interactor. A) Validation assay to test whether 6 of the top interactors identified in the screen can be affinity co-purified with TwinStrep-tagged GPR133-H543R in HEK293T cells. All candidate interactors were tagged with the Myc epitope. Ai) Western blot stained against Myc-tag and GPR133 (antibody against the cytosolic C-terminus) of whole cell lysates. Aii) Western blots of eluates after StrepTactin purification. Note that PTK7, and to a lesser extent TFRC, co-purify with GPR133. A representative blot of three biological repeats is depicted. B) Membrane topology of PTK7. Note that the juxtamembrane portion of PTK7 is cleaved by MT1-MMP (membrane type 1 matrix metalloproteinase) and ADAMs (a disintegrin and metalloproteinases). C) PTK7 is identified as one of the top interactors on the same volcano plot as depicted in Figure 1D. D) Normalized intensities of peptides corresponding to GPR133 and PTK7 as detected in the mass spectrometry analysis in Figure 1E, detailing all biological replicates. E) Comparison of the GPR133 binding interaction of PTK7 vs. the previously reported ligand PLXDC2 using the StrepTactin purification paradigm. PTK7 and PLXDC2 (prey) were tagged with the Myc epitope for detection, while GPR133-H543R (bait) was tagged at the C-terminus with TwinStrep tag for purification. Note that PTK7 co-purifies with GPR133, while PLXDC2 does more co-purify at detectable amounts under the same experimental conditions.

We then compared the PTK7-GPR133 interaction side-by-side with PLXDC2, the recently described GPR133 ligand expressed in the female reproductive system (Bianchi *et al*., 2021). However, we observed that PLXDC2 was neither detected in our initial screen in the context of patient-derived GBM cells, nor did it show detectable co-purification with GPR133 in HEK293T cells in our hands (Fig. 2E).

To further understand the nature of the interaction between GPR133 and PTK7, we conducted structure-function assays. First, we deleted the pentraxin (PTX) domain in the N-terminus of GPR133 and found that it is not necessary for the interaction with PTK7 in co-purification experiments (Fig. 3A; Fig. EV3A; n=2 independent experiments). Second, when we used the His-tagged PTK7 as the bait in co-purification assays, we detected not only the binding of PTK7 to the full-length uncleavable GPR133-H543R, but also to a secreted GPR133 NTF (Fig. 3B). These findings suggest that the NTF of GPR133 mediates the interaction with PTK7, and that, within the NTF, the PTX domain is not necessary for PTK7 binding.

**Figure 3.**
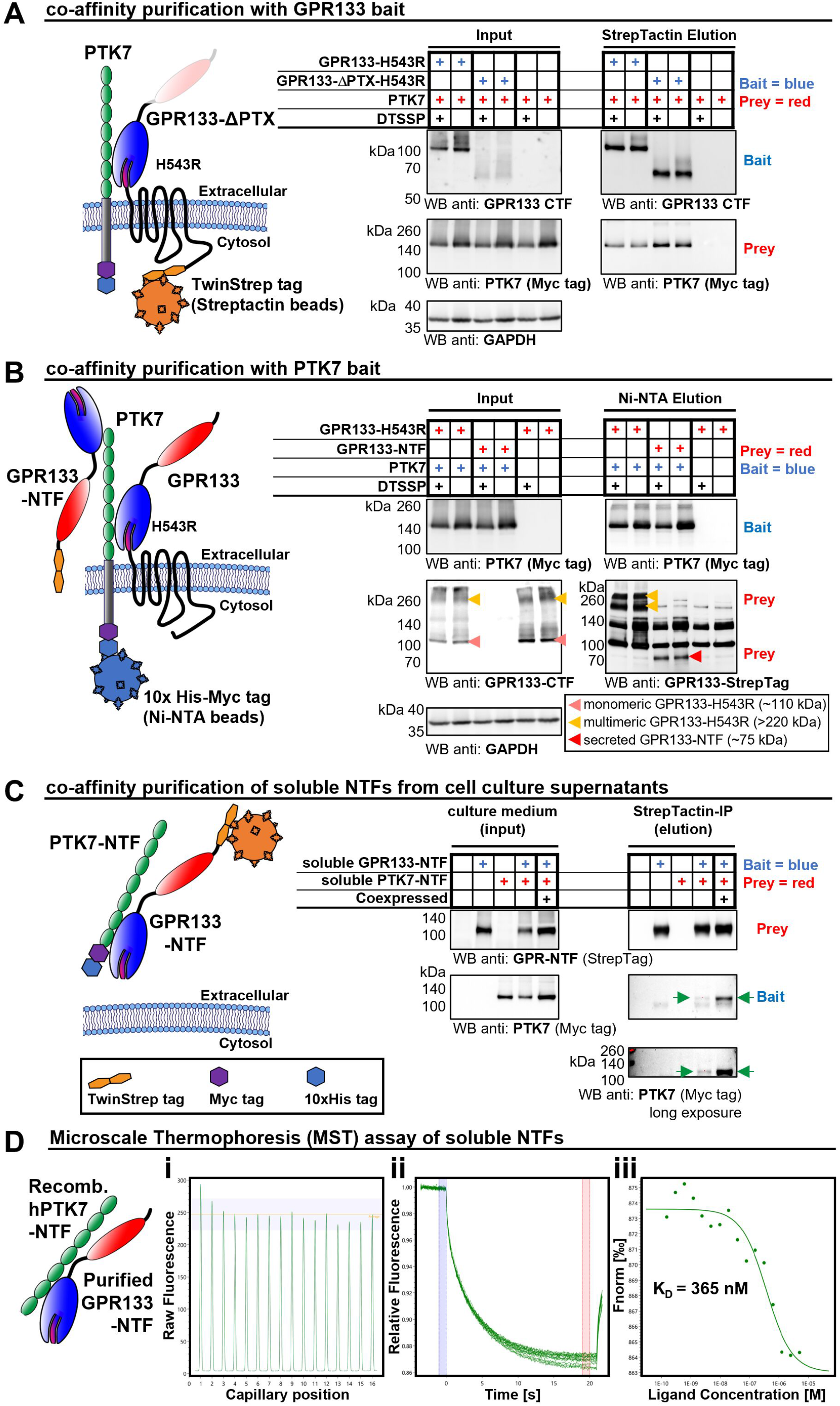
Biochemical characterization of the GPR133-PTK7 interaction. A) Co-affinity purification of different GPR133 variants with coexpressed PTK7. We generated a TwinStrep-tagged GPR133-H543R deletion mutant lacking the PTX domain (GPR133-ϕλPTX-H543R, contains the endogenous signal peptide) and compared its PTK7 binding to full length GPR133-H543R using the StrepTactin purification paradigm. The interaction was not altered by deletion of the PTX domain. Note that inclusion or omission of DTSSP, an extracellular crosslinker, did not influence the binding. Representative Western blots are shown. B) Reverse co-purification experiment using PTK7 as the bait to determine binding to full length GPR133-H543R, or secreted GPR133 NTF tagged at the N-terminus with TwinStrep tag and lacking transmembrane domains. Both the NTF and full length GPR133-H543R co-purified with PTK7 using the Ni-NTA method. Pink arrowheads denote the monomeric full length GPR133-H543R. Yellow arrowheads denote multimers of the full-length GPR133-H543R. Red arrowheads denote the secreted GPR133 NTF lacking transmembrane domains. Multimerization of full length GPR133 due to its transmembrane domains is reduced but not entirely prevented by the use of 1% DDM. This multimerization is discussed in detail in Frenster *et al*. 2021. C) Affinity co-purification of soluble secreted GPR133 NTF and PTK7 NTF. GPR133 NTF with N-terminal TwinStrep tag and secreted Myc-tagged PTK7 with no transmembrane domains were either expressed alone, together in adjacent cells, or co-expressed in the same cells. Secreted PTK7 co-purified with GPR133 in both in mixed-culture and co-expression conditions (green arrows). A representative blot is depicted. D) Quantitative microscale thermophoresis (MST) detects direct GPR133 NTF to PTK7 NTF binding in the absence of other proteins. MST measurements were conducted at 25°C using a fixed recombinant commercial PTK7 NTF concentration (25 nM) and serial dilutions of purified GPR133 NTF starting at 5 µM. (i) A capillary scan was performed prior to the MST experiment to verify negligible fluorescence intensity inhomogeneities between samples and absorption on capillary walls. (ii) MST traces of each sample of the dilution series. No signs of aggregation or ligand-induced photobleaching were detectable. The colored panels highlight the data points used to determine the initial fluorescence (blue) and the fluorescence 20 s after the IR-laser is switched on (red). (iii) Dose-response curve fitted by MO.control, using the “K_D_-model” for analysis. A K_D_ of 365 nM was determined for the interaction of the extracellular fragments of GPR133 and PTK7.

To test directly whether the extracellular NTFs of GPR133 and PTK7 bind each other independently of their transmembrane domains, we expressed secreted extracellular fragments of both proteins in either co-expression or mixed culture systems (Fig. 3C). In co-purification assays, these soluble NTFs co-eluted reproducibly, both in the co-expression and in the mixed culture configurations. The co-purification was stronger when these extracellular fragments were expressed in the same cells, likely due to the increased relative abundance and probability of interaction when the two proteins are co-expressed. Nonetheless, the mixed culture aspect of this assay demonstrated that the extracellular domains of GPR133 and PTK7 bind each other even when expressed on neighboring cells.

Lastly, to obtain a quantifiable measure of the direct interaction between the extracellular fragments of PTK7 and GPR133, we conducted microscale thermophoresis (MST) assays using purified soluble GPR133 NTF (Fig. EV3C, D) and commercially available recombinant human PTK7 NTF. Indeed, we observed that the two receptor NTFs bound to each other with a K_D_ of ∼365 nM in the absence of other proteins, indicating a direct interaction (Fig. 3D).

Taken together, our biochemical assays demonstrate that PTK7 and GPR133 robustly bind each other and co-purify independently of the purification method and the choice of bait. The soluble extracellular NTFs of both proteins are sufficient for binding, even when expressed from different cells in a mixed culture system, but the interaction does not require the N-terminal PTX domain of GPR133. This extracellular PTK7-GPR133 NTF interaction is direct and has a K_D_ of ∼365 nM in MST assays.

### PTK7 expression in *trans* activates GPR133 signaling

GPR133 signals through Gαs-mediated intracellular cAMP increases (Bohnekamp & Schoneberg, 2011). To test whether PTK7 influences GPR133 signaling, we used HEK293T cells, which provide a good experimental system due to their absence of endogenous GPR133 background expression. When co-expressing GPR133 and PTK7 in the same cells, we observed no increase in intracellular cAMP levels as measured by homogenous time resolved fluorescence (HTRF) assays normalized to GPR133 cell surface expression as measured with enzyme-linked immunosorbent assays (ELISA) (Fig. EV4Ai-iii). However, when expressing GPR133 and PTK7 in two separate but co-cultured cell populations, we observed increases in cAMP levels (Fig. EV4Bi-iii). To maximize the interaction surface of these co-cultured cells, we designed a three-layer sandwich system in which GPR133-expressing cells were seeded between two layers of PTK7-expressing cells (Fig. 4A; Fig. EV4D). We used empty vector controls in both cell populations. In this system, expression of PTK7 in the outer layers increased cAMP levels produced by GPR133 in the middle layer (Fig. 4B; Fig. EV4Ci-iii). Importantly, this action of PTK7 depended on its anchoring in the membrane of cells in the outer layers because expression of a secreted PTK7 without its transmembrane domain did not elicit an effect on GPR133 signaling (Fig. 4B). Furthermore, signaling by the cleavage-deficient GPR133-H543R was not influenced by expression of PTK7 in neighboring cells (Fig. 4B). Taken together, these data suggest that GPR133 signaling is increased by PTK7 binding in *trans*, and that both GPR133 cleavage and PTK7 membrane anchoring are required for this activation.

**Figure 4.**
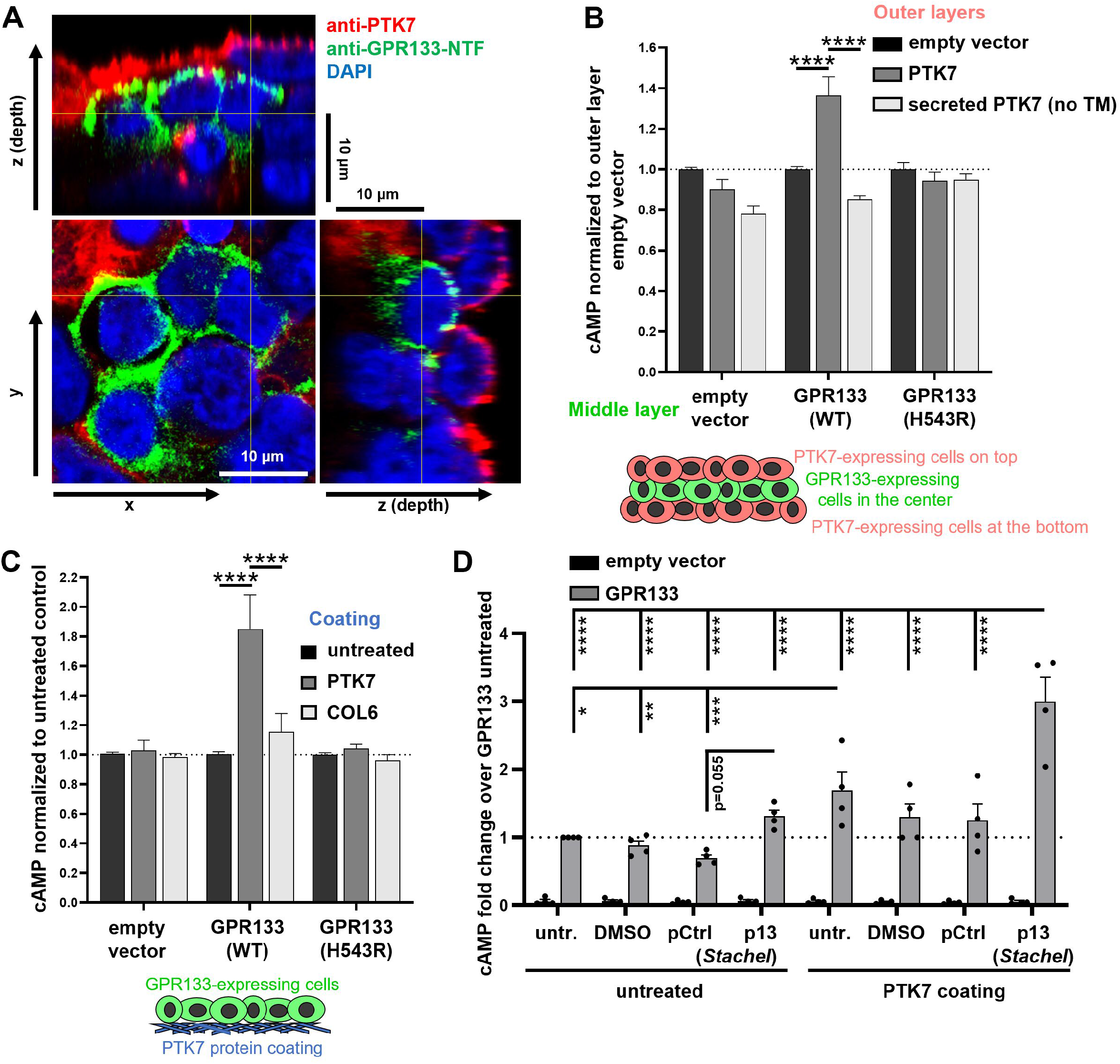
PTK7 increases GPR133 signaling through a *trans* interaction. A) Confocal microscopic image from sandwich cultures shows cells expressing either GPR133 (green) or PTK7 (red). Top view from a central confocal slice (center panel: x and y dimension) and orthogonal views (upper panel: z and x dimension; and right panel: z and y dimension) are shown on the same scale. B) HTRF assays in such sandwich cultures demonstrate that PTK7 in neighboring cells increased cAMP levels in cells expressing wild-type (WT) cleavable GPR133, but not the uncleavable H543R mutant. A secreted version of PTK7 with no transmembrane (no TM) domain did not influence GPR133 signaling (two-way ANOVA: middle layer F_(2, 96)_ = 4.62, not significant; outer layer F_(2, 96)_ = 3.93, not significant; interaction of inner and outer layer F_(4, 96)_ = 7.29, p<0.0001; *post hoc* Tukey’s multiple comparisons: inner layer GPR133-WT expressing cells with empty vector vs PTK7 co-culture, p<0.0001; GPR133-WT expressing cells with PTK7 vs secreted PTK7 (no TM) co-culture, p<0.0001; all other testing not significant; n=4-22 independent experiments). C) Pre-coating culture vessels with PTK7 significantly increased cAMP signaling in WT GPR133-expressing cells, but not cells expressing H543R uncleavable mutant receptor. COL6 (native purified human collagen VI) had no effect on signaling (two-way ANOVA: GPR133 variant expressed F_(2, 80)_ = 11.65, p<0.0001; protein coating F_(2, 80)_ = 12.90, p<0.0001; interaction of factors F_(4, 80)_ = 9.24, p<0.0001; Tukey’s multiple comparisons: GPR133-WT expressing cells on PTK7-coated vs uncoated dishes p<0.0001; GPR133-WT expressing cells on PTK7-coated vs COL6-coated dishes, p<0.0001; all other testing not significant; n=5-15 independent experiments). D) Combinatorial effect of PTK7 binding and p13 *Stachel* peptide treatment on GPR133 signaling. HEK293T cells expressing either WT GPR133 or empty vector control were seeded onto PTK7 NTF-coated or control-coated wells. Cells were then treated with synthetic p13 *Stachel* peptide, inactive control peptide, or solvent controls, and cAMP levels were measured by HTRF. Note how the combination of PTK7-NTF binding and *Stachel* peptide elicited a supralinear response in GPR133 signaling compared to the individual treatments (Two-Way ANOVA, GPR133 expression effect F_(1,48)_=365.1, p<0.0001; treatment effect F_(7,48)_=13.2, p<0.0001; interaction of GPR133 expression and treatment F_(7,48)_=13.2, p<0.0001; Tukey’s multiple comparisons: * = p<0.05; ** = p<0.01; *** = p<0.001; **** = p<0.0001; n=4 independent experiments).

To further support the notion that the extracellular N-terminus of PTK7 needs to be tethered to the plasma membrane or another rigid substrate for this activation mechanism to occur, we used purified human extracellular domain of PTK7 to coat cell culture dishes. Indeed, we observed that GPR133-expressing cells seeded on PTK7-coated dishes exhibited significantly higher cAMP levels when compared to cells cultured on untreated dishes (Fig. 4C; Fig. EV4E). This effect could not be replicated by coating dishes with other proteins, such as collagen VI (COL6, which was detected in a previous unpublished screen for GPR133 interactors by our group, not included in this publication), suggesting that the GPR133-PTK7 interaction is specific, rather than due to promiscuous adhesion. Again, cleavage-deficient H543R mutant GPR133 signaling remained unaffected by PTK7 coating, indicating that the GPR133-PTK7 interaction requires GPR133 cleavage to promote signaling. It is noteworthy that the PTK7 NTF does not increase GPR133 signaling when secreted in a soluble form despite binding GPR133, but does activate GPR133 signaling when rigidly immobilized on cell culture dishes (Fig. 4B and 4C). These findings support a model in which PTK7 binds the GPR133 extracellular domain *in trans*, and via mechanical forces generated by the binding either causes conformational changes and/or facilitates the dissociation of the GPR133 NTF from its CTF, which in turn activate receptor signaling.

To test how the PTK7 binding allosterically influences GPR133’s response to orthosteric agonism by synthetic p13 peptides mimicking the endogenous *Stachel* agonist (Liebscher *et al*., 2014), we performed cAMP HTRF assays in GPR133-expressing HEK293 cells cultured on PTK7-coated wells and treated with p13 *Stachel* peptide (Fig. 4D). While both PTK7 binding and administration of *Stachel* peptide alone increased GPR133 signaling, the combination of both resulted in a supralinear synergistic effect. This suggests that PTK7 binding initiates receptor conformational changes that make the orthosteric *Stachel* binding pocket in the transmembrane portion more accessible. Taken together, our signaling assays point towards a mechanism in which binding of PTK7 to the extracellular domain of GPR133 *in trans* exerts mechanical forces that allosterically increase receptor signaling. The effect may be mediated by conformational changes making the orthosteric *Stachel* binding pocket more accessible. Such conformation changes may also include NTF-CTF dissociation.

### PTK7 and GPR133 have complementary expression patterns in glioblastoma and healthy human tissues

To interrogate whether these PTK7-GPR133 interactions could occur in human tissues, we first investigated the expression patterns of PTK7 and GPR133 within human GBM biospecimens. Epifluorescent and confocal microscopic analysis of surgically resected GBM biospecimens (n=5) revealed that both proteins are expressed throughout tumors (Fig. 5A, B; Fig. EV5A, B;), confirming previous reports of PTK7 being expressed in GBM (Liu *et al*., 2015). We observed spatial gradients in the expression of both proteins, frequently suggestive of a complementary expression pattern, and confirmed in higher magnification confocal Z-stacks that there are indeed adjacent cells predominantly expressing either PTK7 or GPR133 (Fig. 5C). This observation is aligned with our model that it is the *in trans* interaction of GPR133 and PTK7 on adjacent cells that allosterically increases GPR133 signaling.

**Figure 5.**
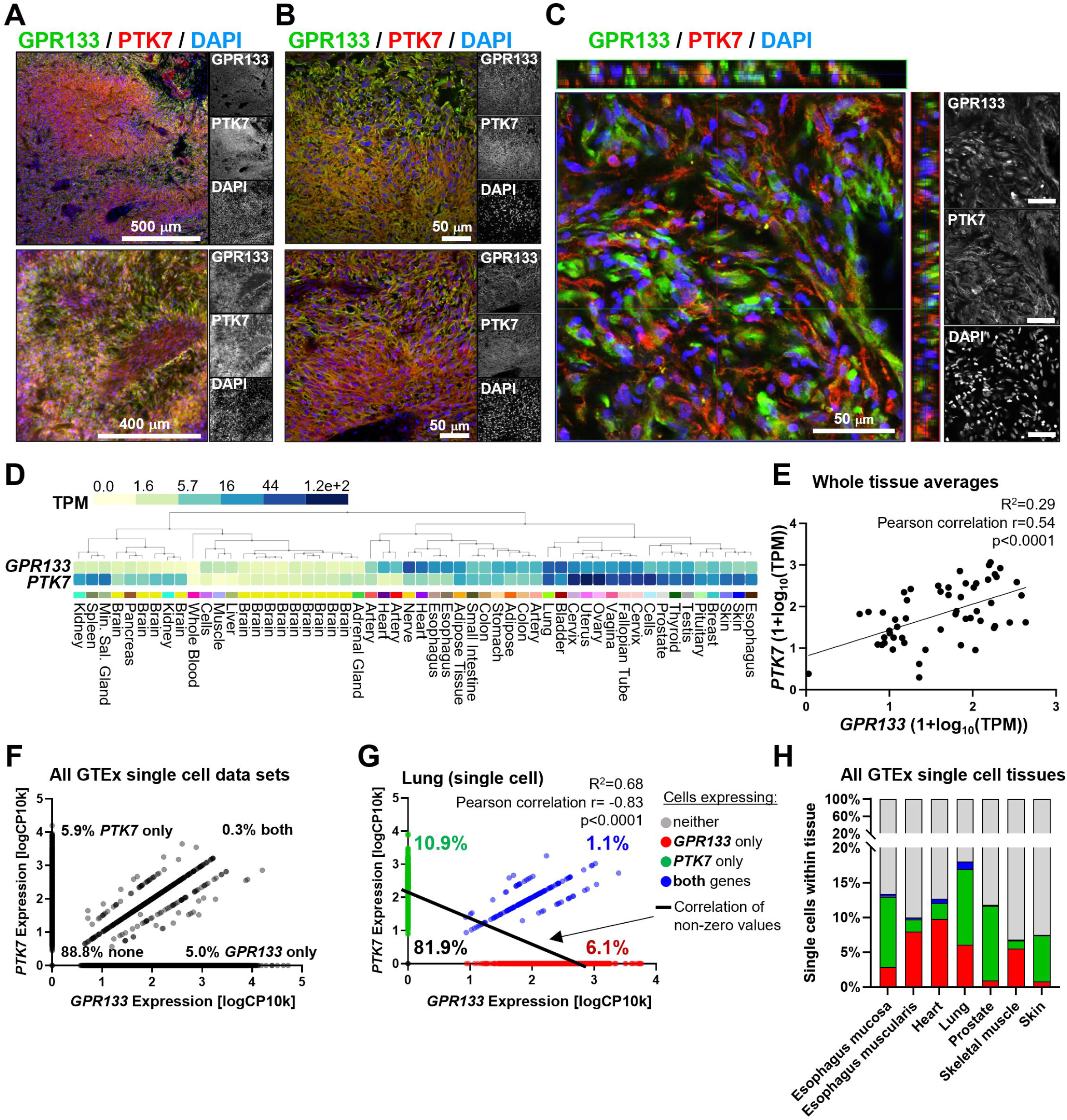
GPR133 and PTK7 expression profiles in GBM and healthy human tissues. A) Epifluorescence microscopic images from two GBM surgical specimens show expression of both GPR133 (green) and PTK7 (red) within tumors. B) Higher magnification epifluorescence images demonstrate a complementary expression pattern in GBM specimens. C) Confocal microscopic image with orthogonal views of a Z-stack indicates that cells tend to either predominantly express GPR133 or PTK7. D) GTEx portal mRNA data from human tissues shows overlap in *PTK7* and *GPR133* expression in several tissues. E) On a whole tissue level, the expression of *PTK7* mRNA showed a significant positive correlation with *GPR133* mRNA. Each dot represents the whole tissue average gene expression from the human GTEx tissues listed in panel D. F) Single cell gene expression analysis of the 7 available GTEx single cell data sets: esophagus mucosa, esophagus muscularis, heart, lung, prostate, skeletal muscle, skin, filtered for cells with RIN>8, demonstrates that *GPR133* only (5%) and *PTK7* only expressing cells (5.9%) are more common that cells co-expressing both genes (0.3%) (130,840 cells included, single cell expression normalized as log10 of copy number per 10,000 reads). G) Within the lung, which exhibits high expression of *GPR133* and *PTK7* on a whole tissue level, individual cells preferentially express only *GPR133* (6.1%) or only *PTK7* (10.9%) rather than co-expressing both genes (1.1%), resulting in a negative correlation (Pearson correlation r = −0.83, R^2^=0.68, P<0.0001, n=2,597 cells; cells expressing neither gene were excluded from correlation analysis to avoid zero inflation). H) All 7 available single cell data sets from the GTEx portal reflect the same negative correlation between *GPR133* and *PTK7* expression in human tissues observed in panel G, with *GPR133* only (red) and *PTK7* only expressing cells (green) being more common that cells expressing both genes (blue). Detailed analyses of these tissues are depicted in Figure EV6.

To gain insight into whether the GPR133-PTK7 interaction may be relevant to physiological processes, we interrogated the GTEx expression database for *GPR133* and *PTK7* mRNA levels across healthy human tissues. Indeed, we found co-expression of the two transcripts in several tissues (Fig. 5D; Fig. EV5C). This co-expression was demonstrated by a positive correlation between *GPR133* and *PTK7* transcript levels across human tissues (Pearson correlation r=0.54, p<0.0001) (Fig. 5E). The specificity of this correlative expression pattern is further supported by the fact that when subjecting all significantly enriched GPR133-binding proteins of this study (n=23) to hierarchical clustering by their similarity of expression across all tissues, *PTK7* and *GPR133* are paired closest to each other (Fig. EV5D), with the exception of GPR89. To determine the cell type-specific expression of *PTK7* and *GPR133* transcripts within these tissues, we interrogated single cell RNA-seq data from the GTEx portal. This analysis indicated that despite the positive correlation on tissue-level, individual cells expressing only *GPR133* or only *PTK7* were more common than cells co-expressing both genes together (Fig. 5F-H). This negative correlation of *GPR133* and *PTK7* expression on a single cell level became even more apparent when looking at single cells within individual tissues (Fig. EV6), exemplified by the lung, which showed high expression for both genes on a whole tissue level (Fig. 5G; Pearson correlation r=-0.83, p<0.0001). This often complementary, inversely correlated expression of the two transcripts in different cell types within the same tissue stands in support of our findings that GPR133 signaling is activated by PTK7 only when they are expressed *in trans*.

### PTK7 knockdown in patient-derived GBM cultures phenocopies GPR133 knockdown

PTK7 was previously implicated in the pathogenesis of GBM (Liu *et al*., 2015). We theorized that, if PTK7 and GPR133 interact in GBM, knockdown of either protein should produce similar phenotypes. In order to test this prediction, we used patient-derived GBM cells and lentiviral shRNA against PTK7 and GPR133. Multiple different shRNA constructs against PTK7 were tested to instill confidence in our findings. Western blot analysis confirmed a reduction in endogenous PTK7 protein levels following knockdown with all tested shRNAs (Fig. 6Ai, ii). Tumorsphere formation assays (Bayin *et al*., 2016) were then used to assess the effect of PTK7 on clonogenic growth potential. Indeed, sphere formation was impaired with multiple knockdown constructs targeting PTK7, similar to the effects of GPR133 knockdown (Fig. 6Bi, ii; tested in 2 independent patient-derived GBM cell cultures), and in accordance with our previous observations (Bayin *et al*., 2016). The fact that knockdown of PTK7 phenocopies the effects of GPR133 knockdown is consistent with our hypothesis that the two proteins interact in GBM.

## DISCUSSION

The long N-termini of adhesion GPCRs are theorized to serve dual roles: first, they adhere to transmembrane proteins or components of the extracellular matrix; and second, they modulate receptor signaling. Here, we demonstrate that PTK7, a type I single-pass transmembrane protein, binds the N-terminus of the aGPCR GPR133 (Fig. 3) to allosterically increase receptor signaling (Fig. 4). The effect is observed only when PTK7 and GPR133 interact *in trans*, i.e. on neighboring cells (Fig. 4B). In addition, the allosteric activating influence of PTK7 on GPR133 signaling requires the membrane insertion of the former and the autoproteolytic cleavage of the latter (Fig. 4B). Furthermore, PTK7 synergizes with soluble *Stachel* peptide to supralinearly activate GPR133 signaling (Fig 4D). Collectively, these data suggest a model in which the PTK7-GPR133 interaction transduces mechanical forces that allosterically induce conformational changes in GPR133. In the PTK7-bound state, the enhanced exposure and accessibility of the orthosteric *Stachel* binding pocket in the transmembrane portion of the receptor leads to increased receptor signaling.

*PTK7* and *GPR133* mRNA expression levels positively correlate across non-pathological human tissues in bulk RNA-seq analyses, but inversely correlate on a single-cell level within these tissues, suggesting their *in trans* interaction is relevant to physiological processes (Fig. 5D-H; EV5C,D; EV6). Importantly, they are both expressed in GBM, a brain malignancy that depends on GPR133 for growth *in vitro* and *in vivo* (Fig. 5A-C) (Bayin *et al*., 2016; Frenster *et al*., 2020). The fact that knockdown of PTK7 phenocopies that of GPR133 in patient-derived GBM cells is consistent with our hypothesis that the PTK7-GPR133 interaction is relevant to GBM pathogenesis (Fig. 6).

**Figure 6.**
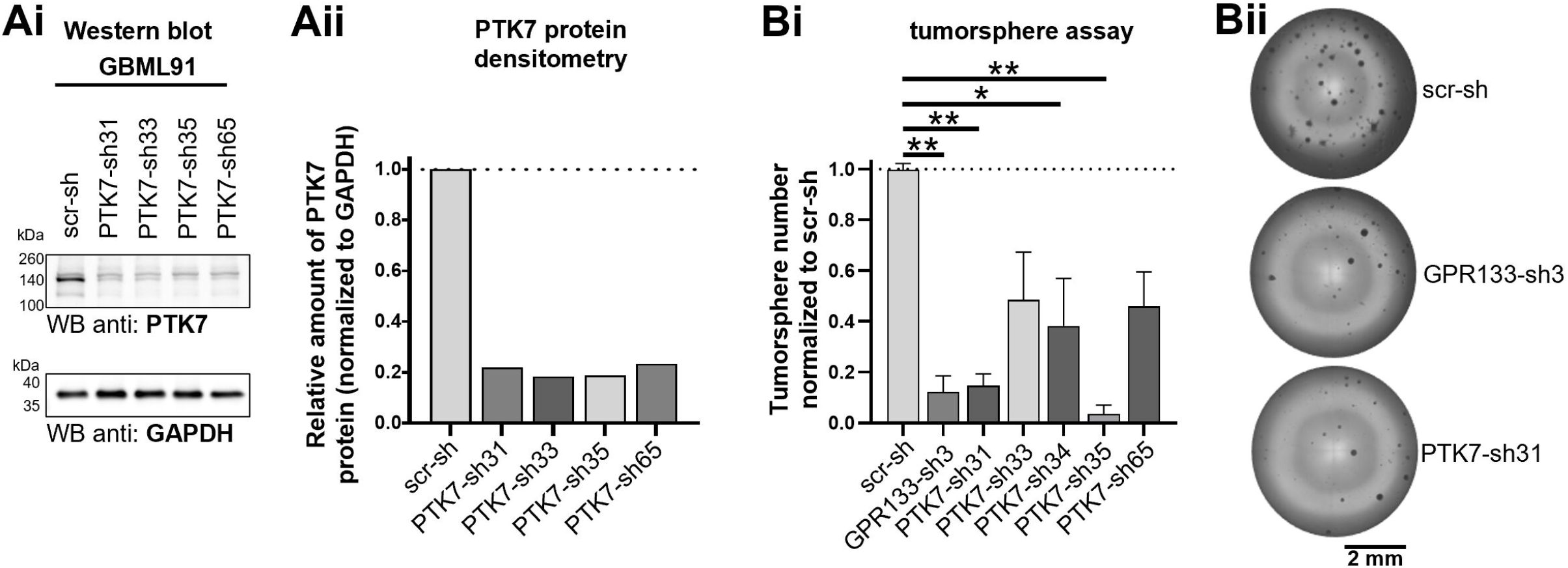
Knockdown of PTK7 phenocopies that of GPR133 in patient-derived GBM cells. A) We tested four different shRNAs targeting PTK7 in patient-derived GBM cells and found significant knockdown for all in Western blots (Ai) and densitometric analysis (Aii). A non-targeting scrambled shRNA (scr-sh) was used as control. Bi) shRNAs targeting PTK7 reduced tumorsphere formation *in vitro*, similar to an shRNA targeting GPR133. Bii) Representative example of tumorsphere formation assay.

Our biochemical analyses indicate that the N-terminal fragment (NTF) of GPR133 is sufficient for the interaction with PTK7 (Fig. 3C, D), with a K_D_ of 365 nM, and that its portion extending from the N-terminus to the end of the PTX domain (aa 31-276) is not required for PTK7 binding (Fig. 3A). These observations suggest that the interaction depends on sequences distal to the PTX domain within the N-terminus of GPR133, possibly including the GAIN domain. Of course, these findings do not exclude the possibility that *in vivo* the interaction may additionally rely on sequences within the extracellular portion of the CTF of GPR133. The *in trans* allosteric effect of PTK7 on GPR133 signaling is abolished when GPR133 is rendered uncleavable by the H543R mutation or when PTK7 is not inserted in the plasma membrane, but rather secreted as a truncated protein lacking its transmembrane and intracellular portions (Fig. 4B). Conversely, the positive allosteric effect of this truncated PTK7 is restored when it is anchored on a rigid surface (Fig. 4C). These findings provide insights into mechanistic aspects of GPR133 activation by PTK7 (Fig. 7). We propose that membrane-tethered PTK7 binds GPR133’s NTF and exerts a mechanical “pull” force that results in conformational changes that may include dissociation of the NTF and the C-terminal fragment (CTF). This new conformation is predicted to allosterically increase GPR133 signaling by facilitating the agonistic interaction of the endogenous *Stachel* sequence with its orthosteric binding pocket within the transmembrane portion of the receptor (Frenster *et al*., 2021; Ping *et al*., 2022; Qu *et al*., 2022). Inherent assumptions in this model are both the intramolecular cleavage of GPR133 that allows for a non-rigid receptor conformation amenable to allosteric modulation, including NTF-CTF dissociation, as well as the membrane insertion of PTK7, which ensures that a mechanical force helps induce the above conformational changes. Consistent with this model is our observation that the allosteric effects of PTK7 on GPR133 signaling synergize with soluble *Stachel* peptide to supralinearly increase receptor signaling (Fig. 4D). In other words, we think that PTK7 binding in *trans* facilitates positioning of the *Stachel* peptide into its binding groove through allosteric effects on protein conformation, whether these include NTF-CTF dissociation or not (Fig. 7).

**Figure 7.**
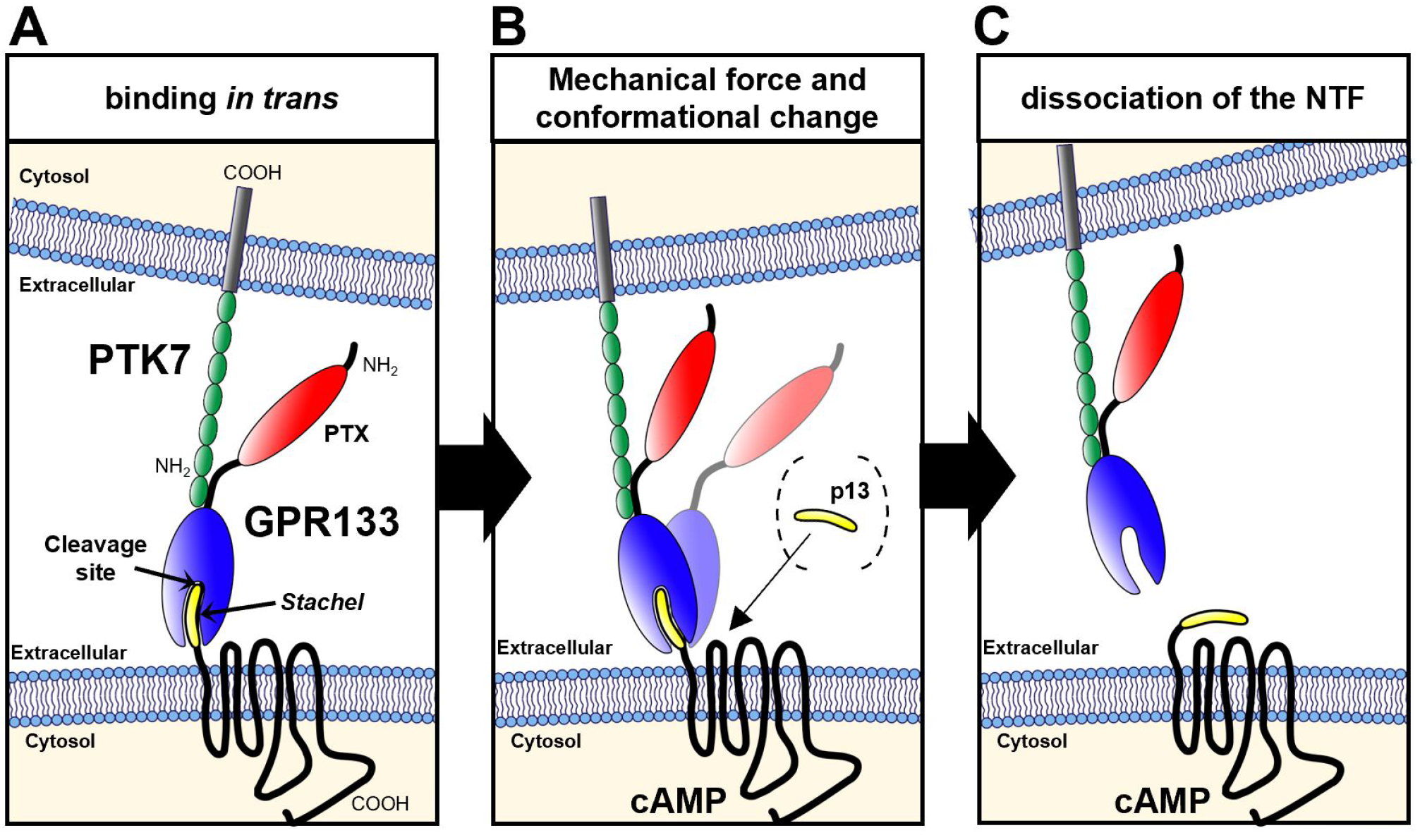
Graphical summary of proposed PTK7-GPR133 interaction model. A) The extracellular portion of PTK7 binds the GPR133 NTF *in trans*. The NTFs are capable of binding each other directly in the absence of other binding partners. B) The binding may induce mechanical forces and a conformational change in GPR133 which makes the orthosteric binding pocket more accessible to the endogenous *Stachel* agonist. This increased accessibility can be detected when adding soluble synthetic p13 *Stachel* peptides to the cells (in dashed brackets). C) Additional mechanical shear stress might facilitate NTF-CTF dissociation, causing the irreversible activation of GPR133 signaling.

The novel interaction between PTK7 and GPR133 will require extensive functional characterization in the future. Our imaging data from GBM operative specimens indicate not only the presence of both proteins in tumors, but also a complementary expression pattern, where cells tend to predominantly, but not exclusively, express one or the other. This profile is consistent with our finding that PTK7 activates GPR133 signaling only when expressed *in trans*, i.e. in adjacent cells, but also raises other important questions. First, our HEK293T data indicate that co-expression of the two proteins in the same cell has no effect on GPR133 signaling. In GBM, tumor cells tend to have a predilection for predominantly expressing one or the other, but overall both seem to be present in the vast majority of cells. A logical future question will be how the relative weights of *cis* and *trans* interactions influence GPR133 signaling. Second, the juxtamembrane extracellular region of PTK7 is subject to proteolysis by metalloproteases, with such cleavages predicted to untether the N-terminus of PTK7 from the membrane. Our experiments have indicated that eliminating the membrane insertion of PTK7 abolishes effects on GPR133 signaling, suggesting that *in vivo* proteolytic cleavage of PTK7 by metalloproteases may also prevent any agonistic effects on GPR133 signaling. Given that metalloproteases are abundantly expressed in GBM, juxtamembrane cleavage of PTK7 represents a potentially crucial regulatory mechanism of the PTK7-GPR133 interaction. Third, we have so far analyzed the agonistic influence of the PTK7-GPR133 interaction on GPR133 signaling, but its effects on PTK7 function are unknown. PTK7 is a known component of Wnt signaling complexes (Dunn & Tolwinski, 2016; Ganier *et al*, 2022; Peradziryi *et al*, 2012), and the Wnt system, in turn, is an important tumorigenic mechanism in GBM (Barzegar Behrooz *et al*, 2022). We, therefore, intend to dissect bidirectional effects of this novel interaction on both PTK7 and GPR133 function in the future.

It is becoming clear that several adhesion GPCRs exhibit promiscuity in interactions through their N-termini. As an example, CD97 (ADGRE5) is known to bind several ligands: CD55, CD90, chondroitin sulfate, and integrin (Hamann *et al*, 1996; Liu *et al*, 2022; Niu *et al*, 2021; Stacey *et al*, 2003; Wandel *et al*, 2012). Similarly, GPR133 was recently shown to interact with PLXDC2, a plexin domain-containing protein, in the female reproductive system. Here, we demonstrate that GPR133 binding to PTK7 in the context of GBM increases receptor signaling. This suggests that the interactome of the N-terminus of GPR133 is context-dependent, and that additional ligands may exist in other tissues where GPR133 is expressed. In our ligand discovery assay, we identified several GPR133 interactors containing Ig domains, such as PTK7, JAM3, NECTIN2, MCAM, NCAM1, and CADM4, although only PTK7 among them showed reproducible binding and modulation of signaling. However, our screen indicated additional interactors lacking Ig domains, for example TFRC. Detailed structure-function studies and functional assays will be needed to elucidate the biochemical basis and phenotypic consequences of such interactions.

In summary, we describe a novel interaction of GPR133 with PTK7, which results in positive allosteric modulation of GPR133 signaling when the two proteins are expressed on adjacent cells. This effect on receptor signaling depends on both the intramolecular cleavage of the N-terminus of GPR133 at the GPS and the membrane insertion of PTK7. Potential mechanisms include PTK7-mediated mechanical forces that allosterically generate conformational changes in GPR133, with or without NTF dissociation, that lead to increased Gs signaling. Additional work will be required to dissect the significance of this interaction in GBM pathogenesis, as well as in physiological processes in tissues where both GPR133 and PTK7 are expressed.

## MATERIALS AND METHODS

### Cell culture

Patient-derived glioblastoma (GBM) cultures were established and maintained as previously described (Bayin *et al*., 2016; Bayin *et al*, 2017; Frenster & Placantonakis, 2018; Frenster *et al*., 2021). In brief, fresh biospecimens from patients undergoing surgical resection of GBM were obtained after informed consent (NYU IRB study 12-01130). These tumor-specimens were mechanically minced with surgical blades and further enzymatically dissociated to single cells using Accutase (Innovative Cell Technologies, Cat# AT104). Patient-derived GBM cells were maintained in spheroid suspension cultures, or if experimentally required, grown as attached cultures on dishes pretreated with poly-L-ornithine (Sigma, Cat# P4957) and laminin (Thermo Fisher, Cat# 23017015). GBM growth medium was formulated as 500 mL Neurobasal medium (Gibco, Cat# 21103049) supplemented with 5 mL N2 (Gibco, Cat# 17-502-049), 500 μL B27 (Gibco, Cat# 12587010), 5 mL non-essential amino acids (Gibco, Cat# 11140050), 5 mL GlutaMax (Gibco, Cat# 35050061), and was freshly supplemented with 20 ng/mL recombinant basic Fibroblast Growth Factor (bFGF; R&D, Cat# 233-FB-01M) and 20 ng/mL Epidermal Growth Factor (EGF; R&D, Cat# 236-EG-01M) every 48 hours. These fresh patient-derived cultures were profiled for mutations and copy number variations (CNV) (Table S1) on an Ion Torrent S5 instrument using a focused next-generation sequencing panel of 50 genes (NYU Oncomine focus assay; Table S2) (Hovelson *et al*, 2015; Mehrotra *et al*, 2017). All parental tumors were isocitrate dehydrogenase (IDH) wild-type GBM.

Lenti-X HEK293T cells (Takara, Cat# 632180) were cultured according to the manufacturer’s protocols in DMEM (Gibco, Cat# 11965-118) supplemented with 10% fetal bovine serum (Peak Serum, Cat# PS-FB2) and sodium pyruvate (Gibco, Cat# 11360070).

All cells were maintained in humidified cell culture incubators at 37°C and 5% CO_2_. HEK293T cells were cultured at 21% O_2_, while patient-derived GBM cells were cultured at 4% O_2_ to more closely resemble physiological conditions.

### Ligand discovery

To detect membrane-bound or extracellular proteins interacting with the full-length GPR133 receptor, three separate patient-derived GBM cultures (GBML61, GBML91, GBML128) were lentivirally transduced with either a C-terminally TwinStrep-tagged and H543R mutated uncleavable GPR133 overexpression cassette, or an empty vector control. Both lentiviral vectors express mCherry for easy identification of transduced cells. Transduced cells were isolated by fluorescence activated cell sorting (FACS) and expanded under adherent culture conditions. For each experiment, ten confluent 10 cm plates for each GBM cell line and construct were used as input and the entire experiment was repeated in three independent experiments. On the day of the experiment, adherent cell cultures were washed once with HBSS (+Ca^2+^, +Mg^2+^) and subsequently fixed using the cell membrane impermeable 3,3’-dithiobis(sulfosuccinimidyl propionate) (DTSSP, Thermo Fisher Scientific, Cat#21578) crosslinking agent at a concentration of 1 mM for 2 hours at 4°C. The crosslinking reaction was quenched by supplementing it with 20 mM Tris. Cells were washed once with ice-cold PBS, scraped off the culture dishes, and pelleted by centrifugation. Cell pellets were lysed in RIPA buffer supplemented with 1% n-dodecyl β-D-maltoside (DDM) (Thermo, Cat# BN2005) and protease inhibitor cocktail (Thermo, Cat# 78429), and with 1/10 volume of 10X Buffer W (IBA, Cat# 2-1003-100) and BioLock (IBA, Cat# 2-0205-250) to block free biotin. After 20 minutes of mixing and incubation on ice, solutions were precleared by centrifugation at 5,000 x g for 15 minutes. The precleared lysates were slowly run through Strep-Tactin sepharose columns (IBA, Cat#2-1202-001) by gravity flow, followed by multiple washing steps, as recommended in the manufacturer’s protocols. All wash buffers were supplemented with 0.1% DDM to remain above the critical micellar concentration of the detergent. The GPR133 bait and all co-immunoprecipitated proteins were eluted using the manufacturer’s elution buffer containing 2.5 mM desthiobiotin (IBA, Cat#2-1000-025). Elutions were analyzed by Western blot and mass spectrometry.

### Protein identification: Nanoflow liquid chromatography-coupled to tandem mass spectrometry (LC-MS/MS)

#### Sample Processing

Affinity purified samples were subjected to SDS/PAGE to fractionate proteins and remove contaminants. The resulting gels were washed 3 times in distilled deionized H_2_O for 15 minutes each and visualized by staining overnight with EZ-Run Protein Gel Staining Solution (Thermo Fisher Scientific, Cat# BP36201). Stained protein gel regions were typically excised into 4 gel sections per gel lane, and destained using standard published protocols (Kishinevsky *et al*, 2018). In-gel digestion was performed overnight with mass spectrometry grade trypsin (Trypsin Gold, Promega, Cat# V5280) at 5 ng/μL in 50 mM NH_4_HCO_3_ digest buffer. After acidification with 10% formic acid (final concentration of 0.5-1% formic acid), resulting peptides were desalted using hand-packed, reversed phase Empore C18 Extraction Disks (3M, Cat#3M2215), following an established method (Rappsilber *et al*, 2007).

#### Mass spectrometry data acquisition

Desalted peptides were concentrated to a very small droplet by vacuum centrifugation and reconstituted in 10 μL 0.1% formic acid in H_2_O. For two technical replicate analyses, ∼90% of the peptide material was used for liquid chromatography, followed by data-dependent acquisition mode (DDA) tandem mass spectrometry (LC-MS/MS). A Q Exactive HF mass spectrometer was coupled directly to an EASY-nLC 1000 (Thermo Fisher Scientific, Cat#LC120) equipped with a self-packed 75 μm x 20-cm reverse phase column (ReproSil-Pur C18, 3M, Dr. Maisch GmbH, Germany) for peptide separation. Analytical column temperature was maintained at 50 °C by a column oven (Sonation GmBH, Germany). Peptides were eluted with a 3-40% acetonitrile gradient over 110 min at a flow rate of 250 nL/min. The mass spectrometer was operated in DDA mode with survey scans acquired at a resolution of 120,000 (at m/z 200) over a scan range of 300-1750 m/z. Up to 15 of the most abundant precursors from the survey scan were selected with an isolation window of 1.6 Th for fragmentation by higher-energy collisional dissociation with normalized collision energy (NCE) of 27. The maximum injection time for the survey and MS/MS scans was 60 ms and the ion target value (AGC) for both scan modes was set to 3e6.

#### Mass Spectrometry Data Processing

The mass spectra files were processed using the MaxQuant proteomics data analysis workflow (version 1.6.0.1) with the Andromeda search engine (Cox *et al*, 2011; Tyanova *et al*, 2016). Raw mass spectrometry files were used to extract peak lists which were searched with the Andromeda search engine against the human proteome and a file containing contaminants, such as human keratins. Trypsin digestion was specified allowing up to 2 missed cleavages with the minimum required peptide length set to be seven amino acids. N-acetylation of protein N-termini, oxidation of methionines and deamidation of asparagines and glutamines were set as variable modifications. For the initial main search, parent peptide masses were allowed mass deviation of up to 20 ppm. Peptide spectral matches and protein identifications were filtered using a target-decoy approach at a false discovery rate of 1%. We used the raw MS1 intensity for protein quantitation.

### Computational analysis of ligand candidates

The resulting list of detected proteins was annotated with subcellular localizations using the “cellular_component” entry of the Ensemble database as reference (Howe *et al*, 2021). Of the 2,138 total detected proteins, 569 with either membrane-, extracellular-, or other non-intracellular annotations were kept. The category “membrane” localization was a combination of the annotation terms: membrane, plasma membrane, integral component of membrane, integral component of plasma membrane, voltage-gated calcium channel complex, apical part of cell, cell junction. The category “extracellular” was a combination of the annotation terms: extracellular exosome, extracellular region, extracellular space. The category “other non-intracellular” was a combination of the annotation terms: cell, cellular_component, endoplasmic reticulum, Golgi apparatus, Golgi membrane, intracellular membrane-bounded organelle, NA. To assess differential protein levels in samples containing the GPR133 bait or the control empty vector, a linear mixed-effects model was built for each protein, with log transformed intensity (log_10_(value+1)) as the response, group membership (GPR133 vs. control) as the fixed effect and GBM culture as the random effect. P values for the fixed effect from all models were then ranked and subjected to multiple testing correction with Benjamini-Hochberg procedure. Mixed effects models were fit using R software and the ‘nlme’ package. Of the correctly localized candidate proteins, only those enriched with the GPR133 bait in all three patient-derived cell lines were kept, resulting in a final list of 87 candidate binding proteins. These 87 candidates were screened for non-specific binding by interrogating the CRAPome database (https://reprint-apms.org/) (Mellacheruvu *et al*, 2013) and were assigned a value from 0 to 716 as a measure of frequency of detection by mass spectrometry in other affinity coimmunoprecipitation experiments.

### Western blot analysis

Whole cell lysates were generated by lysing cells in RIPA buffer (Thermo, Cat#89900) supplemented with Halt complete protease inhibitor cocktail (Thermo, Cat# 78429) and 1% n-dodecyl β-D-maltoside (DDM) (Thermo, Cat# BN2005). After 15 minutes on ice, lysates were sonicated using a Qsonica rod sonicator at 25% power for 3 seconds. Whole cell lysates were precleared by centrifugation at 15,000 x g for 10 minutes at 4°C. Protein concentrations were determined using the DC protein assay kit II (BioRad, Cat# 5000112) and standardized across all samples within an experiment. Protein lysates were reduced in Laemmli buffer (BioRad, Cat# 1610747) containing β-mercaptoethanol at 37°C for 30 minutes, but were not boiled to prevent aggregation of hydrophobic transmembrane regions. Equal amounts of protein were separated by SDS-PAGE and transferred to 0.2 μm nitrocellulose membranes (BioRad, Cat# 1620112). After blocking the membranes in 2% BSA in Tris buffered saline (TBS)-Tween for 1 hour at room temperature, they were incubated with primary antibodies (listed in Table S3) at 4°C overnight and visualized either with HRP-conjugated secondary antibodies for chemiluminescence, or with up to 3 simultaneous fluorescent Alexa Fluor Plus-conjugated secondary antibodies. Images were acquired using the iBrightFL1000 system (Invitrogen).

### Affinity purification with the StrepTactin system

TwinStrep-tagged proteins were affinity purified from whole cell lysates or cell culture supernatants using MagStrep “type3” XT beads (IBA, Cat# 2-4090-002) according to the manufacturer’s protocols. All steps were performed on ice or at 4°C. In brief, whole cell lysates containing 1% n-dodecyl β-D-maltoside (DDM) (Thermo, Cat# BN2005) and protease inhibitor cocktail (Thermo, Cat# 78429) or cell culture medium were pretreated with 1/10 volume of 10X Buffer W (IBA, Cat# 2-1003-100) and BioLock (IBA, Cat# 2-0205-250) for 15 minutes on ice to block free biotin. Solutions were then precleared by centrifugation at 15,000 x g for 10 minutes. After input aliquots were obtained from the precleared protein solutions, MagStrep beads were added and samples were incubated rotating at 4°C overnight. The following day, MagStrep beads were separated using a magnetic rack and washed 5 times in a large excess of 1X Buffer W containing 0.1% DDM and protease inhibitors. Proteins were eluted from MagStrep beads in 3 consecutive steps of 20 μL 2X Buffer BXT (IBA, Cat# 2-1042-025) containing 1% DDM. Elutions were pooled and analyzed by Western blot.

### Affinity purification with His-tag Ni-NTA system

10xHistidine tagged proteins were purified from whole cell lysates using HisPur Ni-NTA beads (Thermo, Cat# 88221), according to the manufacturer’s protocols. All steps were performed on ice or at 4°C. In brief, whole cell lysates containing 1% DDM and protease inhibitor cocktail were precleared by centrifugation at 15,000 x g for 10 minutes and supplemented with 10 mM imidazole (Sigma-Aldrich, Cat# I202). After obtaining input aliquots, pre-washed Ni-NTA beads were added, and samples were rotated at 4°C overnight. The following day, the Ni-NTA beads were pelleted by centrifugation at 700 x g for 3 minutes and were subsequently washed 4 times in PBS containing 50 mM imidazole and 0.1% DDM. Proteins were eluted from Ni-NTA beads in 3 consecutive steps of 20 μL 1 M imidazole in PBS containing 1% DDM. Elutions were pooled and analyzed by Western blot.

### Generation and purification of GPR133-NTF used in microscale thermophoresis assays

A GPR133 ECR construct containing an N-terminal HA and a C-terminal TwinStrep-tag was transiently expressed in HEK293T cells (with the following amino acid sequence: ETGYPYDVPDYAQDHPGFQVLASASHYWPLENVDGIHELQDTTGDIVEGKVNKGIYLKEEKGVTLLYYGRYNSSCISKPEQCGPEGVTFSFFWKTQGEQSRPIPSAYGGQVISNGFKVCSSGGRGSVELYTRDNSMTWEASFSPPGPYWTHVLFTWKSKEGLKVYVNGTLSTSDPSGKVSRDYGESNVNLVIGSEQDQAKCYENGAFDEFIIWERALTPDEIAMYFTAAIGKHALLSSTLPSLFMTSTASPVMPTDAYHPIITNLTEERKTFQSPGVILSYLQNVSLSLPSKSLSEQTALNLTKTFLKAVGEILLLPGWIALSEDSAVVLSLIDTIDTVMGHVSSNLHGSTPQVTVEGSSAMAEFSVAKILPKTVNSSHYRFPAHGQSFIQIPHEAFHRHAWSTVVGLLYHSMHYYLNNIWPAHTKIAEAMHHQDCLLFATSHLISLEVSPPPTLSQNLSGSPLITVHLKHRLTRKQHSEATNSSNRVFVYCAFLDFSSGEGVWSNHGCALTRGNLTYSVCRCTHL|TNFAILMQVVPLELARGHQVALSSIGTDDDDKSAWSHPQFEKGGGSGGGSGGSAWSHPQFEK*). To that end, HEK293T cells were cultured in DMEM supplemented with 10% fetal bovine serum (FBS), 1% GlutaMax, 1% non-essential amino acids and 1% penicillin/streptomycin (all materials obtained from Thermo Fisher Scientific) in T-175 tissue culture flasks (Greiner, growth surface 175 cm2). When confluency was reached, the cells were passaged into roller bottles (CellMaster®, Greiner Bio-One, growth surface 2125 cm^2^). The old medium was removed, and the cells were washed with 20mL DPBS. Then, the cells were incubated with 6mL trypsin-EDTA reagent for 1-2 min. After cell detachment, 14mL growth medium were added to resuspend the cells. Subsequently, cell suspensions were transferred into roller bottles and stocked up to 250 mL with growth medium. Roller bottles were kept in a designated incubator (ShelLab), slowly rolling at 37 °C. The cells were transfected four days after passaging. Here, per roller bottle, 0.50 mg of pHLsec-DNA and 0.1 mg of pAdVantage vector (Promega) were incubated for 20 min with 0.9 mg PEI in 25mL of serum-free medium. During complex formation, the old full medium of the roller bottles was exchanged by 225mL reduced medium, containing only 2% FBS. Finally, the conditioned media were harvested five days after transfection. One mL of BioLock biotin blocking solution (IBA, Cat# 2-0205-250) was added to 1.5 L conditioned medium. The medium was gently stirred for 45–60 min at 4°C, centrifuged at 16,000 x g for 30 min, followed by filtration of the supernatant through a 0.22 μm polyether sulfone membrane (Techno Plastic Products). Prior to affinity chromatography, the filtered conditioned medium was ten-fold concentrated by an ultrafiltration unit (10 kDa molecular weight cut-off (MWCO), GE Healthcare).

All chromatography steps were performed at 4°C using ÄKTAxpress or ÄKTA pure protein purification systems (GE Healthcare). Firstly, the TwinStrep-tagged GPR133 construct was subjected to StrepTactin affinity purification, using a 5 mL StrepTrap HP (GE Healthcare). The washing buffer contained 100 mM Tris pH 8.0 and 150 mM NaCl. The elution buffer additionally contained 2.5 mM D-desthiobiotin. After affinity chromatography, the protein was further purified via size-exclusion chromatography (SEC). A HiLoad® 16/60 Superdex® pg 200 (GE Healthcare) was applied for preparative runs. The eluate of the affinity purification was concentrated to a volume between 1-2 mL before injection using centrifugal filter units (Amicon, Merck millipore) with a MWCO of 10 kDa. The SEC buffer consisted of PBS and 0.1% Tween-20. SEC fractions of the main elution peak were pooled and concentrated as described above to yield a suitable protein concentration for the microscale thermophoresis (MST) assay. Fig. EV3 shows the chromatogram of the SEC run and an SDS-PAGE analysis of the eluted fraction.

### Microscale Thermophoresis (MST)

For conduct microscale thermophoresis, a recombinant extracellular protein construct of human PTK7 (aa 31–704, UniProt #Q13308), containing a C-terminal His6-tag, was purchased from ACROBiosystems. The His6-tag was utilized for fluorescence labeling, employing the Monolith His-Tag Labeling Kit (RED-tris-NTA 2nd Generation kit, NanoTemper). The labeled protein was dissolved in a buffer containing PBS, 0.05% BSA and 0.05% Tween-20. A serial dilution of the purified GPR133-NTF described above was prepared according to the manual of the Monolith His-Tag Labeling Kit. Subsequently, we added 50 nM labeled PTK7 to each sample of the dilution series and thoroughly mixed the samples. MST measurements were conducted using the NanoTemper Monolith NT.115 device at 25°C. The samples were transferred to standard Monolith NT.115 capillaries (NanoTemper) and inserted into the instrument. MST was observed at “high” MST-power and 100% LED power. The software MO.Control (NanoTemper) was employed for system control and data analysis.

### Immunofluorescent staining and microscopy

Fresh surgical GBM biospecimens were obtained after informed consent (NYU IRB study 12-01130) from patients undergoing surgical resection and were immediately fixed in 4% paraformaldehyde (PFA, Sigma-Aldrich, Cat# P6148) solution at 4°C overnight. Specimens were then transferred to 30% sucrose solution and incubated at 4°C for 48 hours, before embedding and freezing in Tissue-Plus OCT (Thermo, Cat#23-730-571). Specimens were cryosectioned into 30 μm-thick slices, recovered on Superfrost glass slides (Thermo, Cat#12-550-15) and stored at −20°C. On the day of staining, slides were brought to room temperature, fixed again in 4% PFA for 10 minutes, washed 3 times with PBS containing 0.1% Triton X-100 (PBS-T) and blocked with 10% bovine serum albumin (BSA) in PBS-T for 1 hour at room temperature. Primary antibodies were mixed in 1% BSA in PBS-T solution at the concentrations indicated in Table S3 and incubated at 4°C overnight. After 3 washes with PBS-T, secondary Alexa fluorophore-conjugated antibodies were added at a concentration of 1:1,000 in 1% BSA in PBS-T and incubated for 1 hour at room temperature. After 3 additional washes in PBS-T, nuclei were counterstained with 500 ng/mL of 4′,6-diamidino-2-phenylindole (DAPI, Sigma, CAT#D8417) for 10 minutes at room temperature. Slides were washed again and mounted under a cover glass. Epifluorescent microscopy was conducted on a Zeiss AxioObserver and confocal laser scanning microscopy was conducted on a Zeiss LSM700. Images were analyzed using ImageJ software.

### Ligand candidate expression plasmids

All expression plasmids are based on the lentiviral vector pLVX-EF1α-mCherry-N1 (Takara, Cat# 631986). Codon-optimized variants of GPR133 were previously published and described (Frenster *et al*., 2021). Ligand candidate cDNAs were obtained from the following commercially available plasmids: PTK7 (Addgene, Cat#65250), PCDHGC3 (SinoBiological, Cat#HG22064-UT), TFRC (Addgene, Cat#69610), NECTIN2 (SinoBiological, Cat#HG10005-G), CADM4 (SinoBiological, Cat#HG16033-G), MCAM (SinoBiological, Cat#HG10115-M), PLXDC2 (SinoBiological, Cat#HG23536-U). All ligand cDNAs were tagged with 10xHistidine and Myc tags and subcloned into the same pLVX backbone to generate bicistronic expression cassettes through a T2A cleavage site (ligand-T2A-mCherry).

### Lentiviral production and infection

Lentiviruses were generated and used as previously described (Frenster *et al*, 2018). In short, Lenti-X HEK293T cells (Takara, Cat# 632180) were grown in 10 cm dishes to 80% confluency and co-transfected with expression plasmids of interest and second-generation packaging plasmids (psPax2 + pMD2.G) using Lipofectamine 2000. Lentiviral particles were collected from the cell culture supernatant at 24, 48, and 72 hours after transfection and concentrated using the Lenti-X concentrator (Contech Takara, Cat# 631231) according to the manufacturer’s instructions. Stably transduced cell lines were generated by lentiviral transduction in the presence of 4 µg/mL protamine sulfate at a multiplicity of infection of three (MOI=3), followed by purification of mCherry-expressing cells via FACS.

### HTRF signaling assays

HTRF assays were performed using the cAMP Gs dynamic kit (CisBio, Cat# 62AM4PEC) following the manufacturer’s protocols with slight modifications (Frenster et al. 2021). In brief, HEK293T were either stably transduced with lentiviruses or acutely transfected with expression plasmids using Lipofectamine 2000 (Invitrogen, Cat# 11668-019), according to the manufacturer’s protocol. Twenty-four hours after transfection, cells were reseeded onto 96-well plates pretreated with poly-L-lysine (Sigma, Cat# P4707) at a density of 75,000 cells per well.

In experiments where culture vessels were pre-coated with proteins, 96-well plates were pre-treated with poly-L-lysine, rinsed with distilled deionized H_2_O, and coated with 30 μL of 12.5μg/mL PTK7 (R&D, Cat# 9799-TK-050) or COL6 (Rockland, Cat# 009-001-108), resulting in a total of 375 ng of protein per well. Plates were left to air-dry overnight and rinsed with distilled deionized H_2_0 once before plating cells.

Alternatively, cells were plated as multi-layer sandwich cultures by seeding layers of alternating transgenic cells on top of each other with 12 hours’ time intervals between each layer to allow for attachment.

Twenty-four hours after the final seeding of cells, the medium was exchanged with 50 μL of fresh medium with 1 mM 3-isobutyl-1-methylxanthine (IBMX) (Sigma-Aldrich, Cat# I7018-100MG), and cells were incubated at 37°C for an additional 30 minutes. Cells were then lysed and cAMP levels were measured using the cAMP Gs dynamic kit on the FlexStation 3 (Molecular Devices), according to the manufacturer’s protocol.

### Enzyme-linked immunosorbent assay (ELISA)

Cells were seeded on 96-well plates simultaneously with each corresponding HTRF signaling assay, as described above. Twenty-four hours after seeding, cells were washed with HBSS (+Ca^2+^/+Mg^2+^) and fixed with 4% PFA for 20 minutes at room temperature. Whole cell ELISAs were performed under permeabilizing conditions in the presence of 0.1% Triton X-100 throughout, while surface ELISAs were performed under non-permeabilizing conditions in the absence of any detergent. The fixed cells were first blocked in HEK293T medium containing 10% FBS for 1 hour at room temperature, and then incubated with primary antibodies diluted in HEK293T medium at concentrations indicated in Table S3 for 1 hour at room temperature. After 3 thorough washes with PBS, cells were incubated with horseradish peroxidase-conjugated secondary antibodies diluted 1:500 in HEK293T medium for 1 hour at room temperature. After 3 additional washes with PBS, cells were overlayed with TMB-stabilized chromogen for 10 minutes (Thermo Fisher, Cat# SB02) followed by an equal volume of acidic stop solution (Thermo Fisher, Cat# SB04). Absorbance was read at 450 nm using a Synergy H1 multi-mode plate reader (Biotek). Non-specific background was determined using “secondary-antibody only” control wells and subtracted from sample wells.

### *In silico* analysis of tissue expression profiles

Hierarchical clustering of mRNA levels of ligand candidates against *GPR133* by human tissue was conducted using the Multi Gene query of the Genotype-Tissue Expression (GTEx) web portal (https://gtexportal.org/home/multiGeneQueryPage), without exclusion of any available tissues. Single cell transcriptome analysis was performed using the GTEx single cell data set as released by Eraslan et al. (https://www.biorxiv.org/content/10.1101/2021.07.19.452954v1.full; doi: https://doi.org/10.1101/2021.07.19.452954). The raw data file GTEx_8_tissues_snRNAseq_atlas_071421.public_obs.h5ad was extracted using python 2.7.16. Expression values in log(TP10K+1) were combined with the annotation of all cells in R 4.2.0. Single cell expression data for *GPR133* and *PTK7* from 209,126 cells was extracted from the cited dataset and only cells with RIN>8 were used in the analysis.

### Tumorsphere formation assays

Patient-derived GBM cultures were grown as described above. Cells were dissociated to a single cell suspension and infected at a multiplicity of infection of 3 (MOI=3) with lentiviruses expressing shRNAs in the backbone vector pLKO. Cells received either a non-targeting control shRNA (CCTAAGGTTAAGTCGCCCTCG), a previously described shRNA against GPR133 (CCTGCAGGGACTGTTCATATT) (Bayin, *et al*. 2016), or PTK7-targeting shRNAs from the Sigma MISSION library (Millipore Sigma, Cat# SHCLNG-NM_002821): sh31, CACAGGGTTAATGAGTCTCTT; sh33, CCACAGCACAAGTGATAAGAT; sh34, CCTCATGTTCTACTGCAAGAA; sh35, CCTGAGGATTTCCAAGAGCAA; sh65, GCCACTCATCTGCCAACTTTG). Two days after infection, cells were dissociated to single cell suspensions and selected in 5 µg/mL puromycin for 5 days. Seven days post infection, cells were dissociated to single cells, counted, and plated at 500 cells per well (96-well plate) in 100 μL medium. Cells were allowed to grow and initiate tumorspheres in 96-well plates for 2 weeks, while growth factors and fresh media were supplemented every other day. After 2 weeks of growth, the 96-well plates were imaged using automated tile scanning on an EVOS imaging system (Thermo Fisher Scientific). Tile scans of each well were exported, and sphere number was analyzed in ImageJ. An area of 4,000 μm^2^ (sphere diameter ∼70 μm) was used as a minimum size cutoff during analysis. Each experimental condition was quantified from 6 technical replicate wells, which was counted as a single data point in statistical analyses.

### Statistical analysis

Statistical comparisons were conducted in GraphPad Prism (v9). Replicates were represented as mean ± SEM (standard error of the mean), unless otherwise noted. Statistical significance was calculated using Students t-test; and one-way or two-way analysis of variance (ANOVA) with *post hoc* Tukey’s or Sidak’s multiple comparisons tests. GTEx expression correlation was calculated as Pearson r correlation with two-tailed testing. The threshold of significance was set at p<0.05 throughout.

## ACKNOWLEDGEMENTS

We thank the Microscopy core facility at NYU Grossman School of Medicine for assistance with confocal microscopy imaging. We thank Dr. Andreas Kloetgen for helping with the annotation of subcellular localizations of candidate proteins detected during the ligand discovery affinity screen.

## AUTHOR CONTRIBUTIONS (written in CRediT format)

Conceptualization, J.D.F. and D.G.P.; Investigation, J.D.F., H.E.B.; Methodology, J.D.F.; Formal Analysis, J.D.F. and D.G.P.; Resources, D.G.P., N.S., T.A.N., I.L., and T.S.; Writing – Original Draft, J.D.F. and D.G.P.; Writing - Review & Editing, J.D.F., H.E.B., W.L., G.S., N.R.B., D.B., J.W., S.H., B.K., C.W., N.S., I.L., T.S., D.F., T.N., and D.G.P.; Visualization, J.D.F.; Funding Acquisition, J.D.F., G.S., N.R.B., I.L., T.S., T.N., D.G.P.; Supervision, D.G.P.

## EXPANDED VIEW

**Figure 1 Expanded View.**
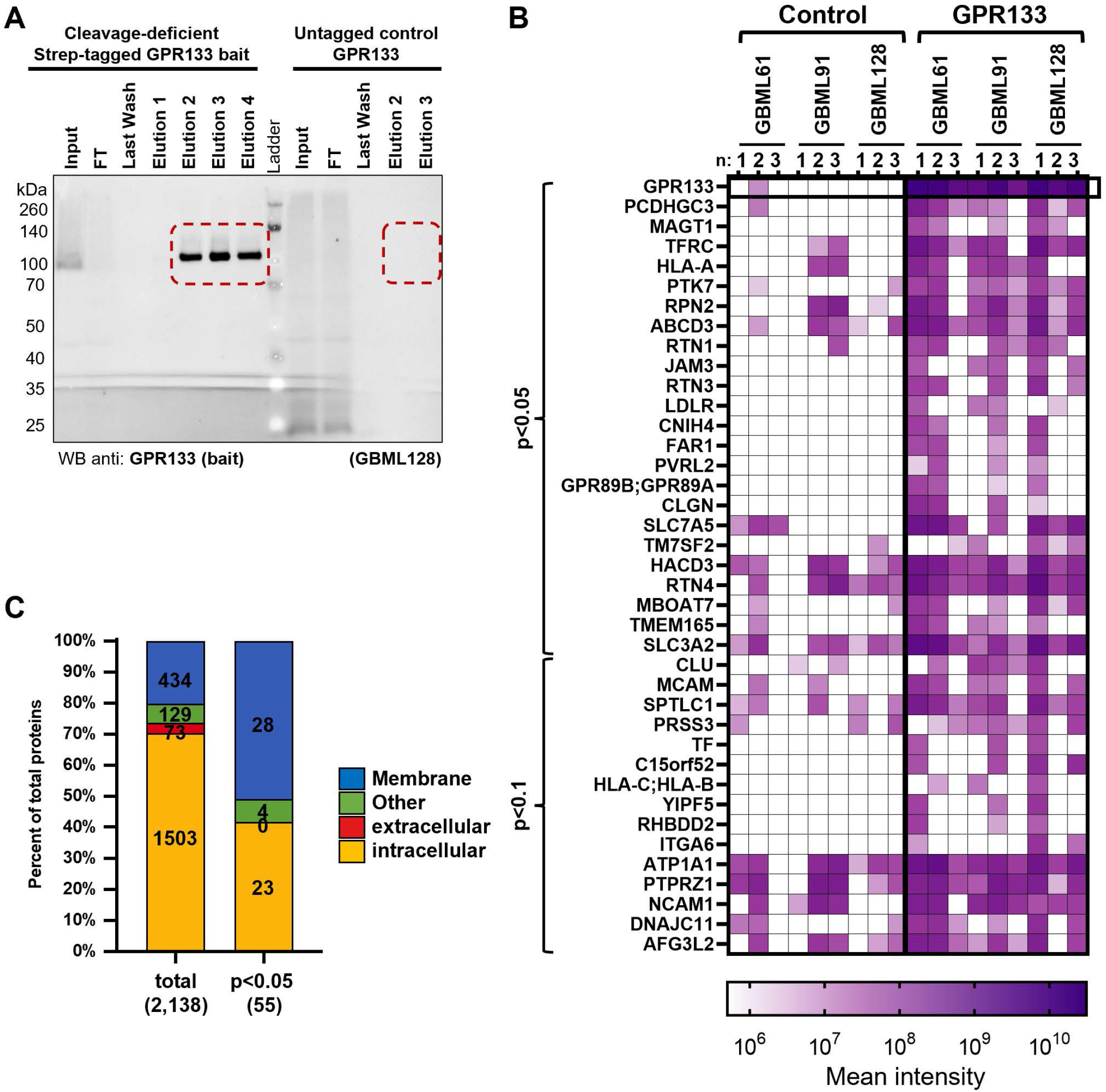
Affinity co-purification with GPR133 bait in patient-derived GBM cells primarily enriches for plasma membrane localized proteins. A) Quality control for the affinity purification of TwinStrep-tagged GPR133-H543R from GBML128 whole cell lysates using StrepTactin columns. Aliquots of the input, flow-through (FT), the last wash, and sequential elution steps were analyzed by SDS-PAGE and Western blot using an anti-GPR133 C-terminus antibody. Tagged GPR133 was strongly enriched in elutions 2-4 compared to the input, while flow-through and wash aliquots did not contain detectable GPR133. Untagged wild-type GPR133 was used as a control (right part of membrane) and was detected in the input and flow-through samples, but not the elutions. B) Heatmap of the 38 top enriched proteins with appropriate subcellular localization (p<0.1) depicting the log_10_ protein intensities of each biological replicate separately. C) Distribution of subcellular localization annotations among the detected proteins from the ligand discovery assay. Plasma membrane-localized proteins are more prominently represented among those proteins enriched with the GPR133 bait (right bar) when compared to the total pool of detected proteins (left bar).

**Figure 2 Expanded View.**
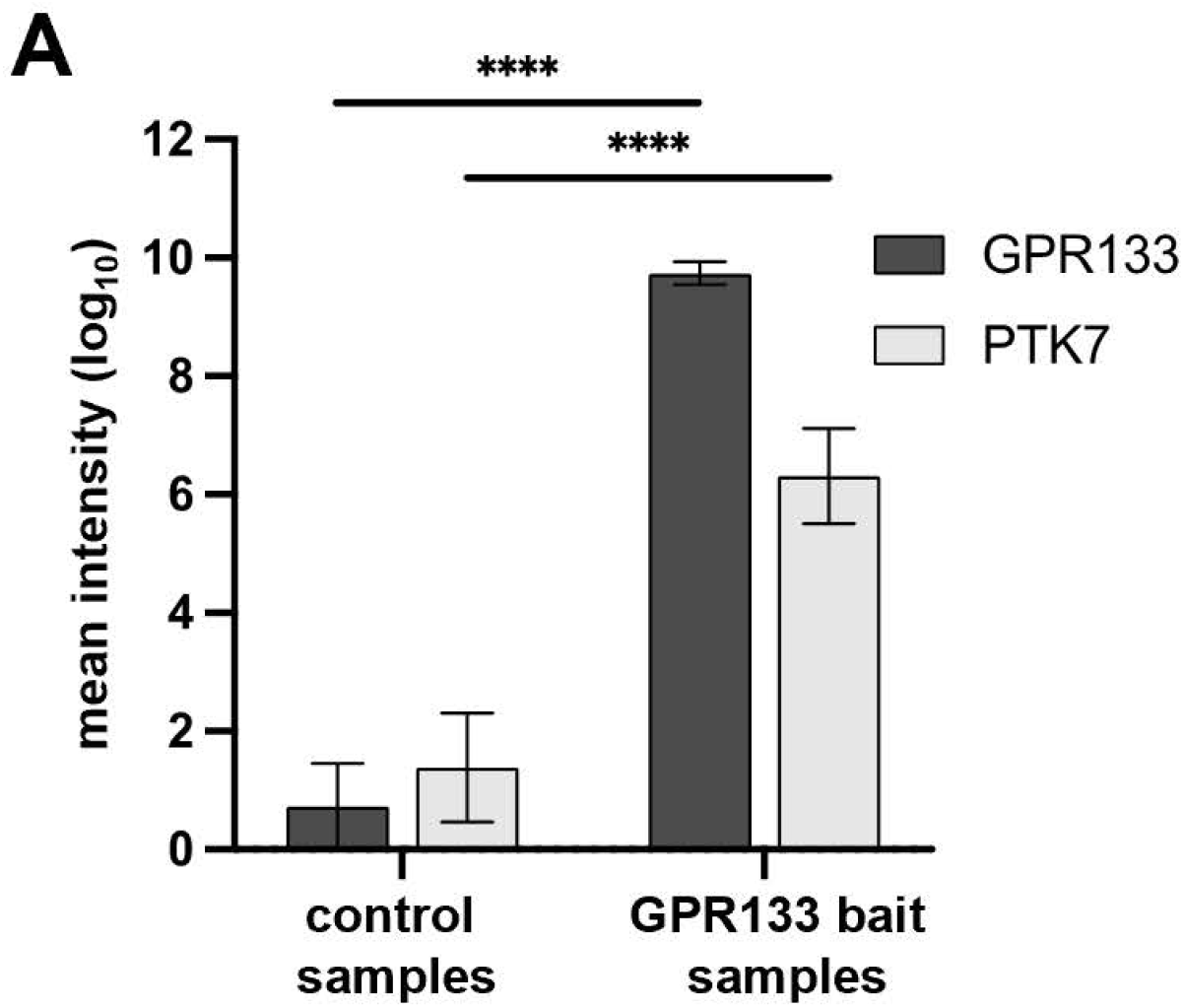
Enrichment of PTK7 with the GPR133 bait in patient-derived GBM cultures. A) Mean intensities (log_10_) +/-SEM of peptides corresponding to GPR133 and PTK7 in the mass spectrometry analysis of the screen. Both GPR133 and PTK7 are significantly enriched in the samples expressing GPR133-H543R (n=3 biological replicates per culture in 3 separate patient-derived cultures; two-way ANOVA GPR133 bait vs. control F_(1, 32)_ = 94.4, p<0.0001; GPR133 vs. PTK7 detection in elution F_(1, 32)_ = 3.71, not significant; interaction of the factors F_(1, 32)_ = 8.12, p<0.01; Sidak multiple comparisons: detected GPR133 protein intensities in control vs. GPR133 bait, p<0.0001; detected PTK7 protein intensities in control vs GPR133 bait, p<0.0001).

**Figure 3 Expanded View.**
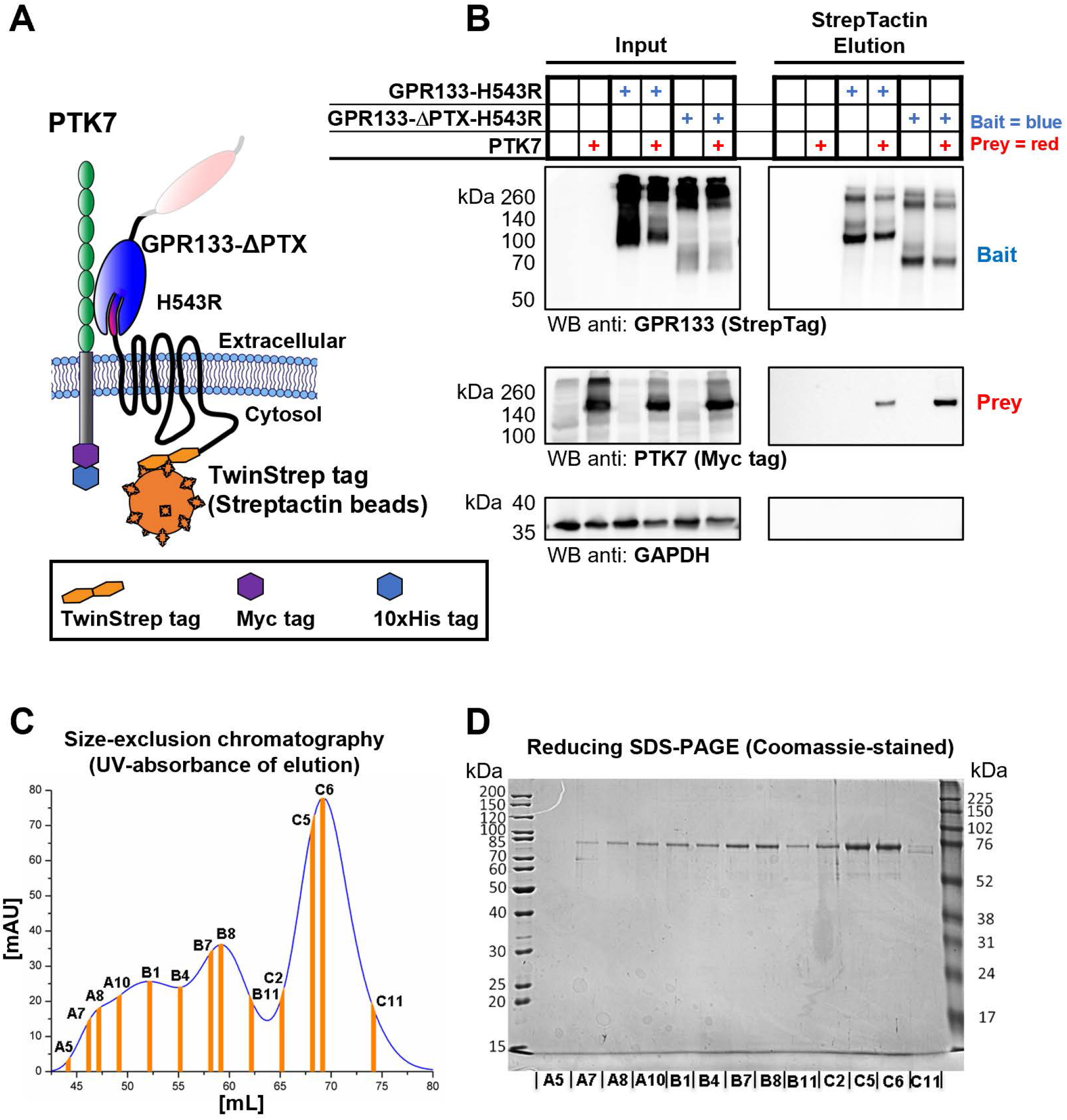
The pentraxin domain is not required for GPR133-PTK7 interaction. A) Schematic showing the co-purification of PTK7 from whole cell lysates using a TwinStrep-tagged H543R mutant GPR133 as bait (GPR133-H543R). B) Western blot analysis of input and elution samples. Removal of the PTX domain within the GPR133 bait (GPR133-ΔPTX-H543R) does not impair the GPR133-PTK7 interaction. The GPR133 bait was detected using an anti-StrepTag antibody, while PTK7 was detected by staining for its intracellular Myc tag. C) SEC chromatogram after GPR133-NTF purification. Fractions taken for SDS-PAGE analysis are highlighted. D) Corresponding Coomassie-stained SDS-PAGE of purified GPR133-NTF demonstrates high degree of purity. Fractions of the main peak were pooled and concentrated for MST assays.

**Figure 4 Expanded View.**
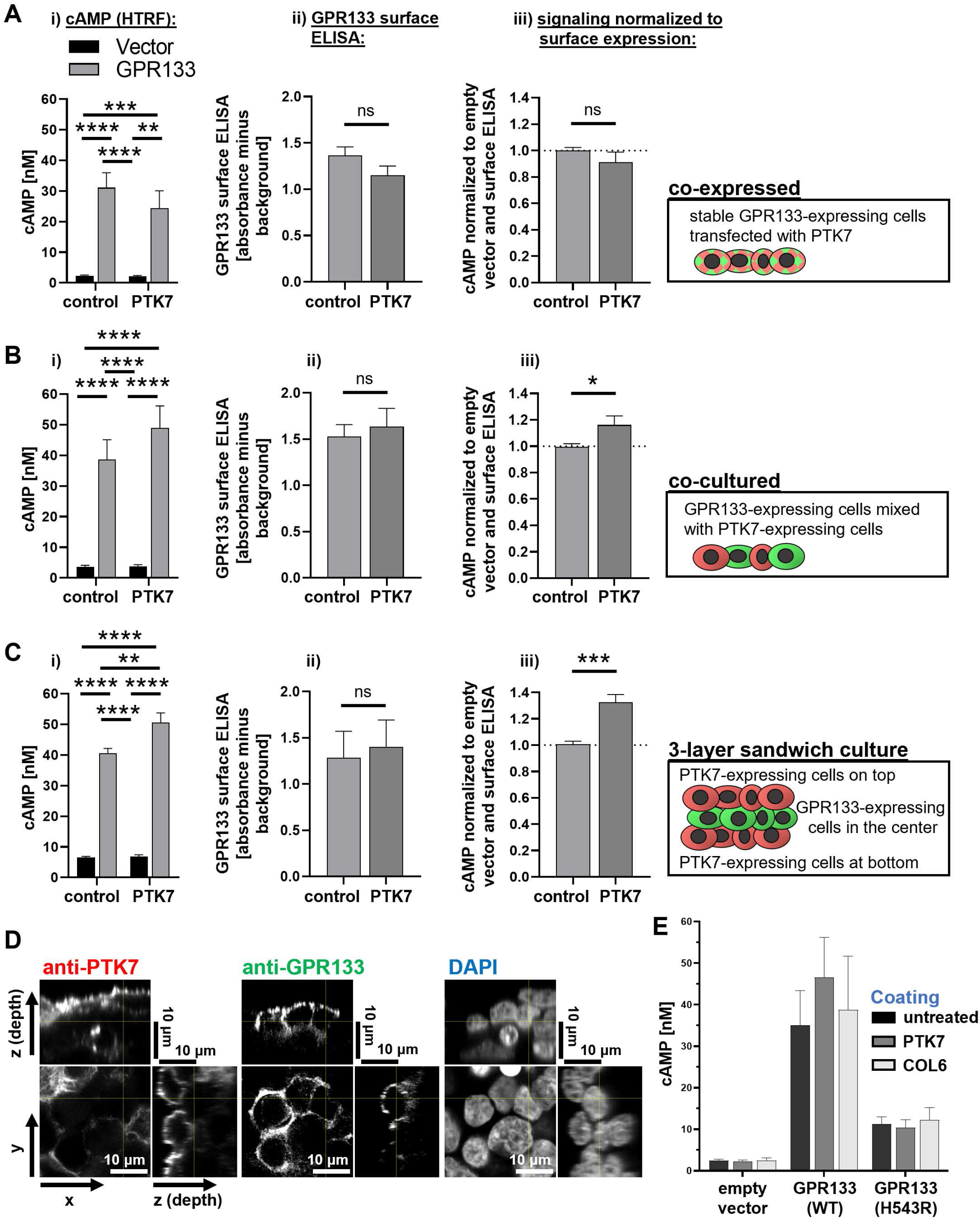
PTK7 interaction in *trans* increases GPR133 signaling. A) GPR133 expressing HEK293T cells or vector control cells were exposed to PTK7 by co-expression via transfection. Ai) cAMP levels were measured by HTRF assays (two-way ANOVA with *post hoc* Tukey’s multiple comparisons: *, p<0.05; **, p<0.01; ***, p<0.001; ****, p<0.0001, n=15). Aii) Surface expression of GPR133 measured by non-permeabilizing ELISA expressed as mean absorbance minus background (unpaired t-tests, not significant, n=13). Aiii) cAMP levels of GPR133 and PTK7 co-expressing cells normalized to GPR133 surface expression (unpaired two-tailed t-test, not significant, n=10 independent experiments). B) GPR133 expressing HEK293T cells or vector control cells were exposed to PTK7 by monolayer co-culture with PTK7-expressing cells in a salt-and-pepper pattern. Bi) cAMP levels were measured by HTRF assays (two-way ANOVA with *post hoc* Tukey’s multiple comparisons: *, p<0.05; **, p<0.01; ***, p<0.001; ****, p<0.0001, n=10). Bii) Surface expression of GPR133 measured by non-permeabilizing ELISA expressed as mean absorbance minus background (unpaired t-tests, not significant, n=11). Biii) cAMP levels of GPR133 expressing cells in PTK7 co-cultures normalized to GPR133 surface expression (unpaired two-tailed t-test, p<0.05, n=7 independent experiments). C) GPR133 expressing HEK293T cells or vector control cells were exposed to PTK7 by creating 3-layer sandwich cultures containing GPR133-expressing cells in the middle layer, surrounded by PTK7-expressing cells at the bottom and top layers. Ci) cAMP levels were measured by HTRF assays (two-way ANOVA with *post hoc* Tukey’s multiple comparisons: *, p<0.05; **, p<0.01; ***, p<0.001; ****, p<0.0001, n=6). Cii) Surface expression of GPR133 measured by non-permeabilizing ELISA expressed as mean absorbance minus background (unpaired t-tests, not significant, n=5). Ciii) cAMP levels of GPR133 expressing cells in 3-dimensional sandwich cultures normalized to GPR133 surface expression (unpaired two-tailed t-test, p<0.001, n=3 independent experiments). D) Corresponding single-color channels of confocal Z-stack microscopy depicted in Figure 4A. E) Absolute cAMP levels of cells expressing an empty vector control, WT GPR133, or H543R mutant GPR133 on either untreated, PTK7-coated, or COL6-coated culture dishes (corresponding to Figure 4C).

**Figure 5 Expanded View.**
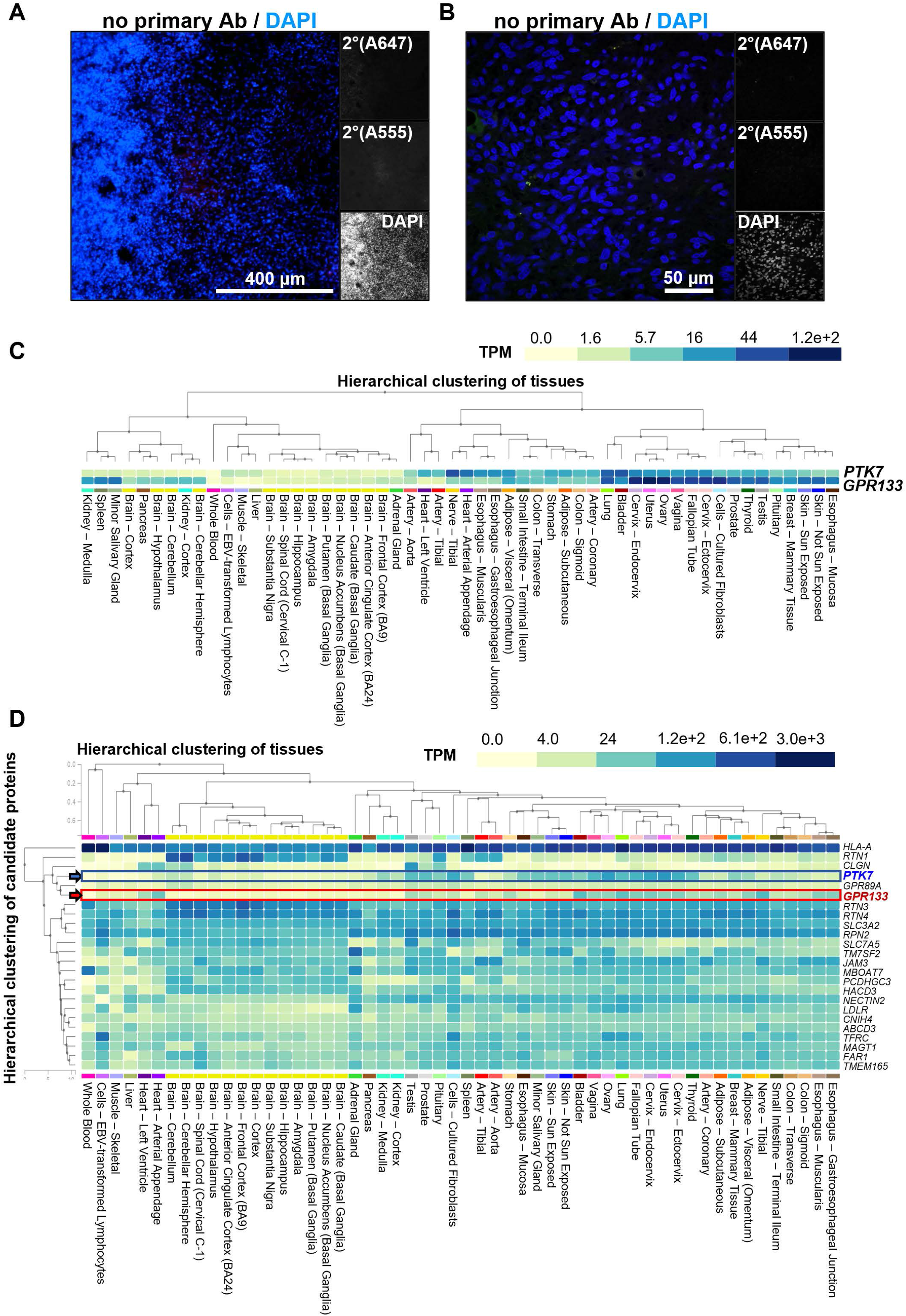
GPR133 and PTK7 expression in GBM and across healthy human tissues. A) Secondary antibody only control immunostaining of operative GBM specimens depicted in Figure 5A. B) Secondary antibody only control stain for the confocal microscopy images of the operative GBM specimen depicted in Figure 5B. Nuclei are counterstained with DAPI. Exposures of all channels were identical to images of Figure 5. C) GTEx multi-gene query of *GPR133* and *PTK7* as depicted in Figure 5, including more extensive tissue labeling. D) GTEx multi-gene mRNA query of the 23 top enriched proteins detected in the ligand discovery screen of Figure 1. Human tissues are hierarchically clustered by the expression profile of these 23 genes. The 23 genes are hierarchically clustered by their similarity of expression across all tissues. Note that *PTK7* (blue) and *GPR133* (red) are clustered closest to each other among all candidate transcripts, with the exception of GPR89.

**Figure 6 Expanded View.**
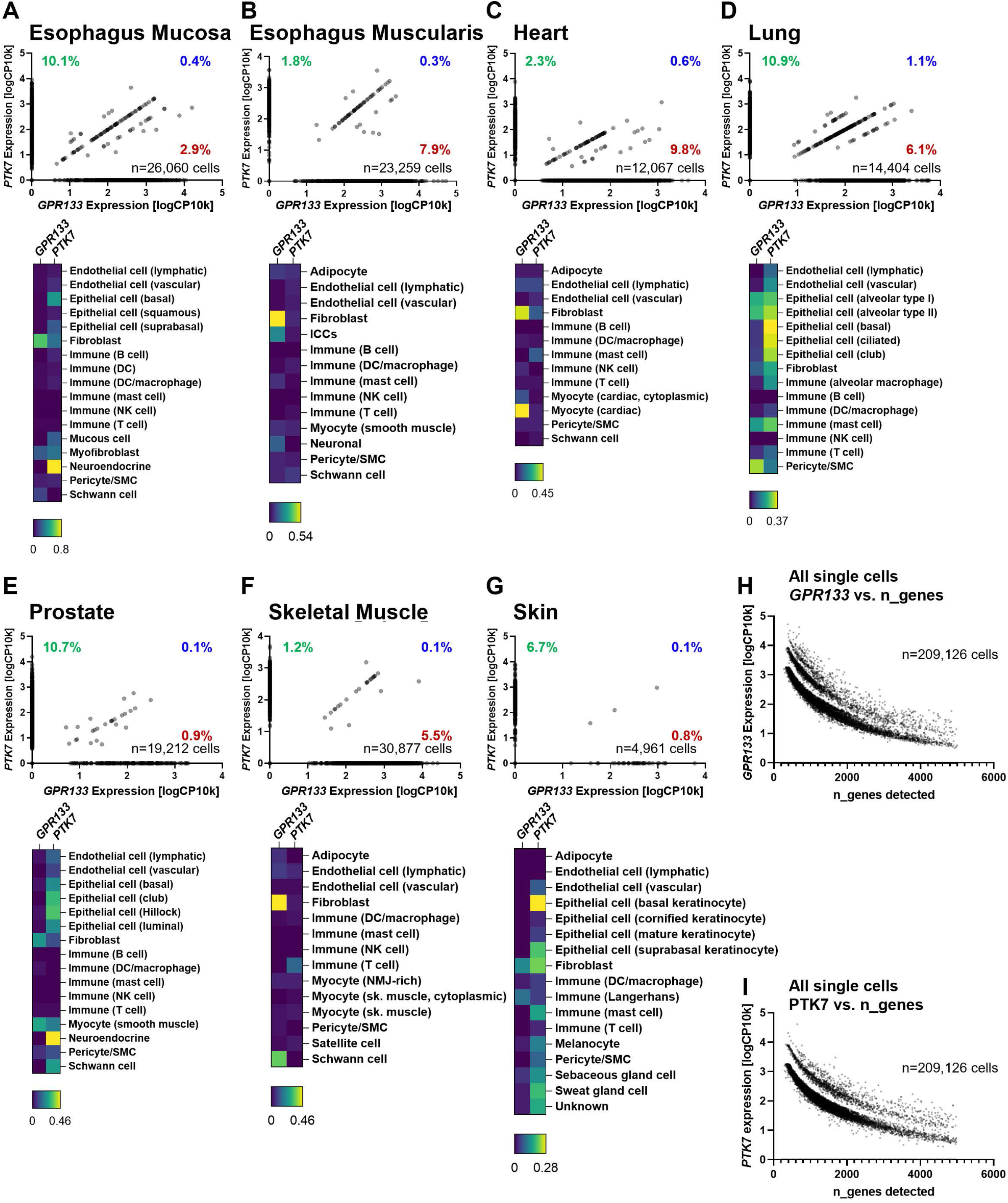
PTK7 and GPR133 are expressed in separate cell populations within healthy human tissues. GTEx single cell RNAseq data was filtered for cells with RIN score >8 and analyzed for the expression of *GPR133* and *PTK7* across the physiological human tissues **A**) esophagus mucosa, B) esophagus muscularis, **C**) heart, **D**) lung, **E**) prostate, **F**) skeletal muscle, and **G**) skin. Dot plots depict single cell expression normalized as log10 of copy number per 10,000 reads. Across all tissues, the percentages of cells expressing only *PTK7* (green) or only *GPR133* (red) are higher than the percentage of cells co-expressing both genes (blue). Heatmaps depict the expression of *GPR133* and *PTK7* within each tissue separated by broad cell types. **H**) *GPR133* expression of all 209,126 available GTEx single cell transcriptomes plotted against the total number of genes detected within a given cell. This quality control confirms that *GPR133* expression is well detected in cells both with overall high gene counts and low gene counts. **I**) *PTK7* expression of all 209,126 available GTEx single cell transcriptomes plotted against the total number of genes detected within a given cell. This quality control confirms that *PTK7* expression is well detected in cells both with overall high gene counts and low gene counts. Together, panels H and I instill confidence in the fact that the detected negative correlation of gene expression is based on biological expression patterns and not artifacts due to low detectability of these genes or low gene coverage. For this analysis only, cells were purposefully not pre-filtered for RIN>8.

## SUPPLEMENTARY TABLES

**Table S1.** Genetic background of patient-derived GBM cultures

| Patient-derived culture | Gender | IDH status | MGMT promoter methylation | EGFR amplification | Oncomine focus assay |
| --- | --- | --- | --- | --- | --- |
| GBML61 | F | wild-type | not methylated | amplified | FGFR3 F386L mutation, JAK3 V722I mutation |
| GBML91 | F | wild-type | methylated | not amplified | no mutations |
| GBML128 | F | wild-type | methylated | not tested | no mutations |

IDH, isocitrate dehydrogenase; MGMT, O<sup>6</sup> methylguanine DNA methyltransferase; EGFR, epidermal growth factor receptor.

**Table S2.** Genes sequenced in NYU Oncomine focus assay

|  |  |
| --- | --- |
| **Mutations** | AKT1, ALK, AR, BRAF, CDK4, CTNNB1, DDR2, EGFR, ERBB2, ERBB3, ERBB4, ESR1, FGFR2, FGFR3, GNA11, GNAQ, HRAS, IDH1, IDH2, JAK1, JAK2, JAK3, KIT, KRAS, MET, MTOR, NRAS, PDGFRA, PIK3CA, RAF1, RET, ROS1, SMO |
| **Copy number variation** | ALK, AR, BRAF, CCDN1, CDK4, CDK6, EGFR, ERBB2, FGFR1, FGFR2, FGFR3, FGFR4, KIT, KRAS, MET, MYC, MYCN, PDGFRA, PIK3CA |
| **Fusion drivers** | ABL1, AKT3, ALK, AXL, BRAF, ERG, ETV1, ETV4, ETV5, EGFR, ERBB2, FGFR1, FGFR2, FGFR3, MET, NTRK1, NTRK2, NTRK3, PDGFRA, PPARG, RAF1, RET, ROS1 |

**Table S3.**
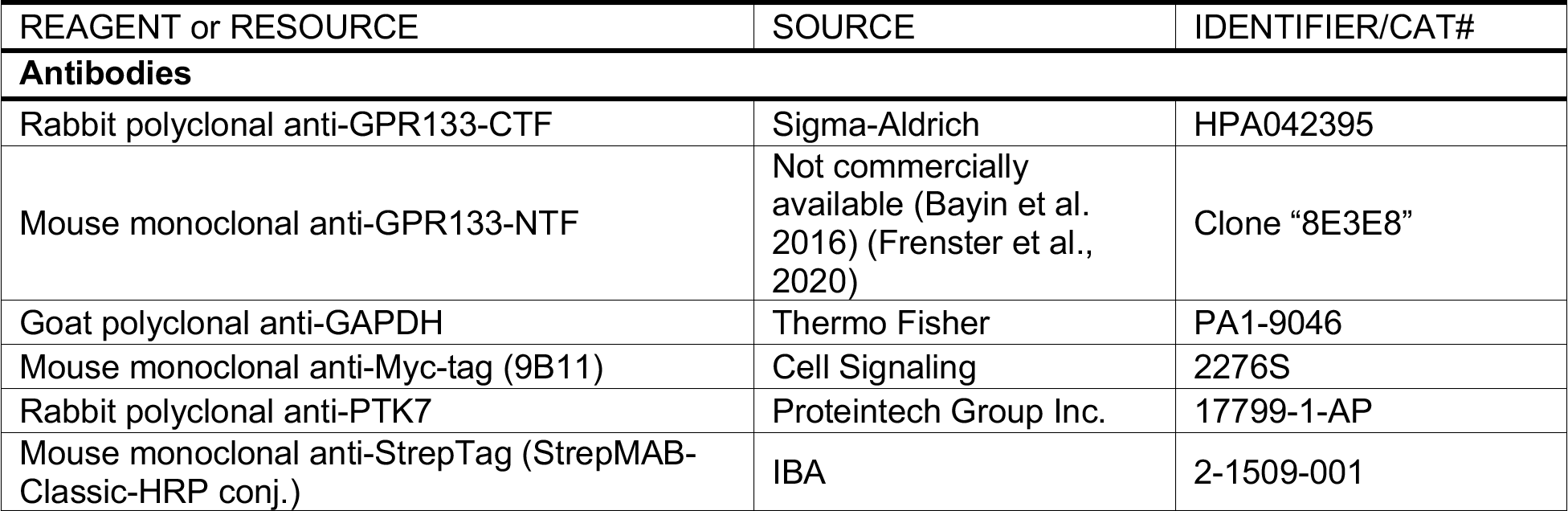
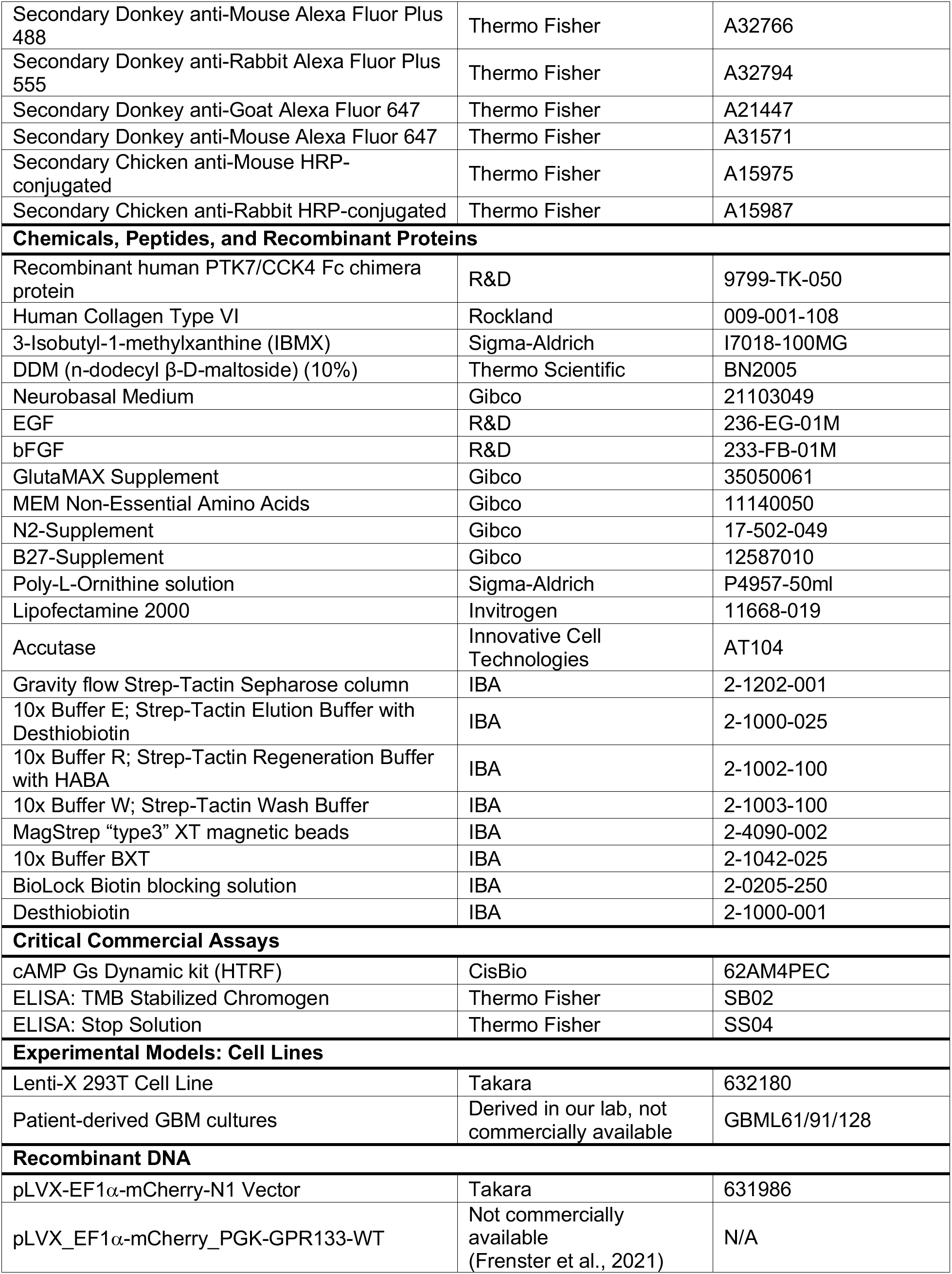
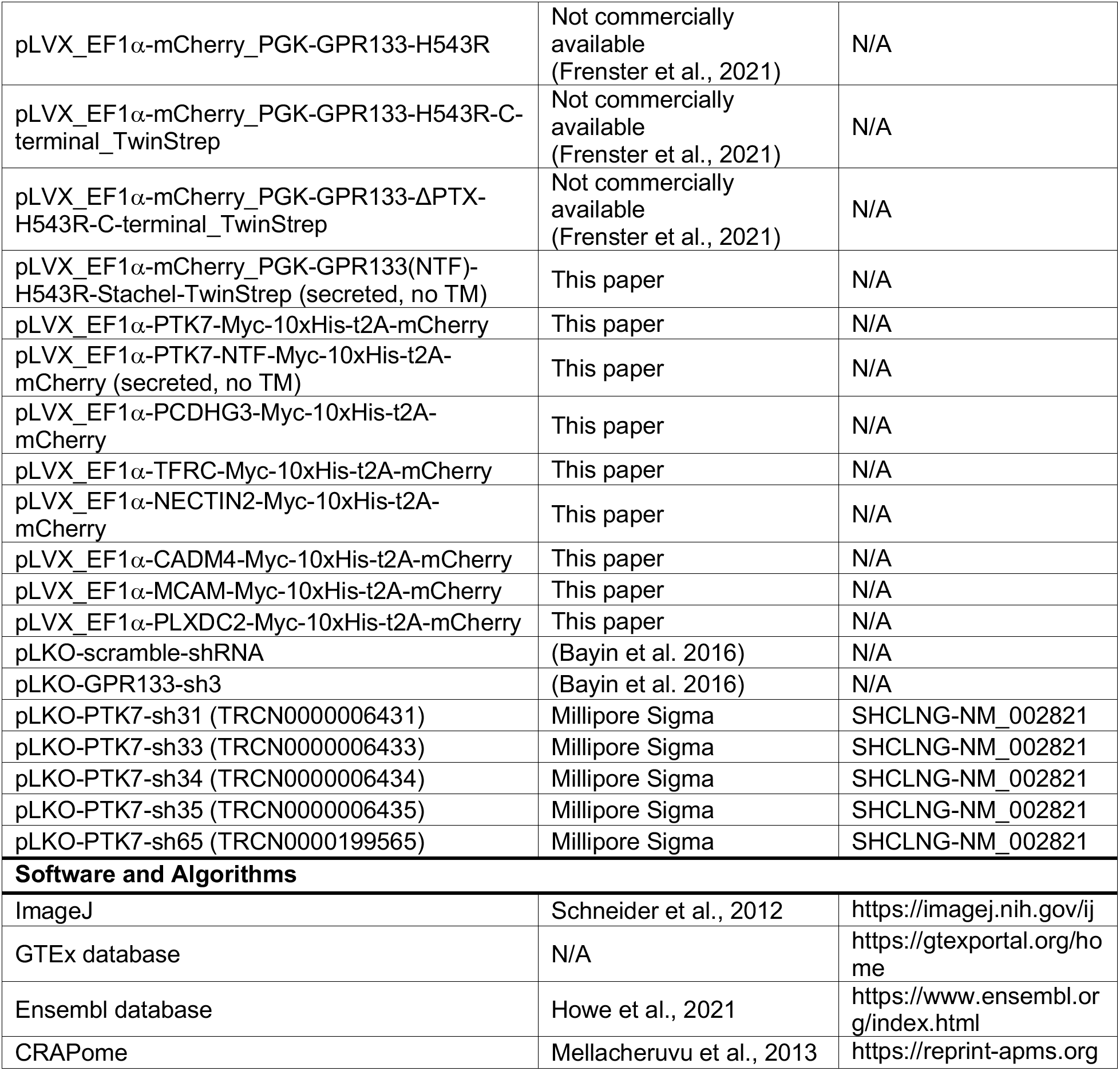
Key reagents

